# A Müller glia-retinal pigment epithelium apical network surrounding cone outer segments in fish and mouse

**DOI:** 10.64898/2026.04.23.719722

**Authors:** Maria Sharkova, Michael Housset, Samantha Boudreau, Chavy Dworkind, Sara Amidian, Michel Cayouette, Jennifer C. Hocking

## Abstract

Homeostasis of the vertebrate retina critically depends on two non-neuronal cell populations, Müller glia and the retinal pigment epithelium (RPE). Their opposing apical surfaces delineate the subretinal space, a specialized extracellular compartment housing the light-detecting outer segments of photoreceptors. Although the individual interactions between these supporting cells and photoreceptors have been extensively characterized, Müller glia and the RPE have long been presumed to remain physically separate in the healthy retina. Here, we challenge this assumption and present a revised model of the subretinal space architecture based on detailed FIB-SEM and confocal microscopy imaging of the zebrafish and mouse outer retina. We demonstrate that both Müller glia and the RPE extend long apical processes that not only contact one another, but also collectively ensheath cone outer segments, forming a previously unrecognized tripartite interface. These contact regions are highly elaborated in fish and, while comparatively sparser, are still present in mouse. Developmentally, the growth and interdigitation of Müller glial and RPE apical processes follow the emergence of the outer segment in both species and, in mouse, coincides with the onset of visual function. Although glial–RPE interactions are enriched at cone-associated sites, their presence is not strictly dependent on cone localization in either species, suggesting they represent a fundamental feature of the vertebrate subretinal space. Together, these findings revise our understanding of the subretinal space cytoarchitecture and reveal a previously unappreciated structural interaction between Müller glia, RPE and photoreceptors that is conserved across vertebrate species.

## Introduction

Vision begins with the capture of light by photoreceptor cells, specialized sensory neurons with a strikingly polarized morphology. Photoreceptors comprise rods, which mediate vision in dim light, and cones, which operate in bright light conditions and support colour vision through spectrally distinct subtypes. The photoreceptor apical domain is subdivided into an inner segment, enriched in organelles, and an outer segment, a cilium-based sensory ending composed of a stack of membranous discs (rods) or lamellae (cones) densely packed with photopigment. Maintaining outer segment architecture imposes substantial energetic and biosynthetic demand, in part because the outer segment undergoes lifelong renewal [1–3]. While the intracellular pathways that build and maintain the outer segment have been dissected in detail, the surrounding extracellular niche remains less well defined.

Photoreceptor outer segments reside in the subretinal space, an isolated and controlled environment positioned between the neural retina and the retinal pigment epithelium (RPE). The subretinal space is delineated proximally by the outer limiting membrane (OLM), formed by cell-cell junctions between Müller glia and photoreceptors, and distally by the tight junctions of the RPE (see schematic depiction in Figure S1A) [4–6]. The importance of the subretinal space is emphasized clinically: when the sensory retina separates from the RPE, as in retinal detachment, disruption of this interface rapidly compromises photoreceptor viability and visual recovery unless hastily corrected [7, 8]. Accordingly, subretinal homeostasis emerges from the coordinated interactions among three major cellular partners: photoreceptors, RPE, and Müller glia [9–11].

RPE cells extend apical processes into the subretinal space, interdigitating with photoreceptor outer segments and contributing to metabolite exchange, retinoid handling and removal of distal outer segment material. The interaction between the RPE and outer segments is best characterized for rods, in part because the individualized rod disc architecture enables visualization of outer segment renewal [1, 2]. In contrast, the RPE–cone interface is less understood than its rod counterpart and varies significantly by species. RPE projections range from microvillar or villous processes to elaborate concentric lamellar sheaths [12–15], sometimes presenting with highly distinct morphologies in comparison to the RPE–rod interface. Notably, the RPE can extend well beyond the tip of the cone outer segment, potentially supporting cone-specific morphogenesis and function within the subretinal space [13, 14, 16, 17].

Müller glia add a second interface to the subretinal space, but the organization and functional significance of their apical domain remain less clearly established. As radial macroglia stretching across the three retinal layers, they provide structural and metabolic support to all retinal neurons [18]. At the apical surface of the neural retina, Müller glia form the OLM through heterotypic specialized junctions with photoreceptor inner segments, creating a mechanically robust boundary that mediates partial diffusion restrictions [6]. Müller glia are frequently depicted as terminating at the OLM with only short apical microvilli extending further. However, ultrastructural and molecular studies across species demonstrate examples of taller actin-based glial processes that reach into the subretinal space, and in some species, run alongside photoreceptor inner segments [18–21]. Still, the structural basis and domains of glial apical processes, their three-dimensional (3D) geometry, and their positioning relative to cones and RPE processes remain poorly defined.

Notably, both Müller glia and RPE apical processes were described extensively in classic ultrastructural work, but largely from flat sections and predominantly in species where genetic manipulation is limited. A comparative, 3D framework is missing in genetically manipulable systems such as zebrafish and mouse, where defining the native architecture is the essential first step before genetics can be used to interrogate mechanism and function.

We previously reported extensive interdigitation of Müller glia and RPE processes around ultraviolet-sensitive (UVS) cones in juvenile zebrafish [17]. Here, we extend our characterization of these subretinal support structures and demonstrate their conservation in mice. Using Focused Ion Beam Scanning Electron Microscopy (FIB-SEM), we generated high-resolution 3D reconstructions of outer retinal cellular relationships, revealing a consistent arrangement of Müller glia and RPE contacts around cones in both species. Notably, the removal of UVS cones did not prevent the formation of these glial–RPE contacts. Together, our analyses establish a cross-species structural framework for interactions between Müller glia and the RPE, providing a basis for investigating how the subretinal architecture supports cone homeostasis and renewal.

## Methods

### Animal husbandry

#### Zebrafish

Zebrafish were housed in a Tecniplast recirculating aquatics system located within the North Campus Animal Services facility of the University of Alberta and maintained in accordance with Canadian Council on Animal Care zebrafish guidelines (ethics protocol: AUP1476). Following breeding, embryos were raised in 10 cm Petri dishes containing embryo medium [1x E2, Zebrafish International Resource Center (ZIRC)] in an incubator set at 28.5°C with a 14/10 light/dark cycle. At 5–6 dpf, larvae were brought back into the facility. Zebrafish were euthanized in an overdose of methanesulfonate salt (Acros Organics, pH 7.0).

The AB strain of zebrafish were used as wild type. Transgenic lines were used to visualize UVS cones [*Tg(sws1:GFP)*] [22], Müller glia [*Tg(gfap:GFP)*] [23], and tagged actin expression in the heat shock experiment [*Tg(hsp:act-myc)*] [17]. *Crystal* zebrafish lacking pigment were a gift from Dr. Ted Allison and generated as previously shown [24, 25]. For the heat shock experiment, young adult (3.5 mpf) zebrafish were kept at 39°C for 1 hour as previously reported [17]. Two hours after heat shock, control and heat shocked groups were further split into two groups each, with one kept in the light (light-adapted), and the other dark-adapted. We euthanized all four groups two hours later; dark-adapted fish were handled under red light.

#### Mice

All mouse work was carried out in accordance with the Canadian Council on Animal Care guidelines and approved by the IRCM Animal Care Committee. Wild-type CD1(ICR) mice were obtained from Charles River Laboratories and used for all RPE electroporation experiments. C57BL/6J mice were obtained from The Jackson Laboratory and used for FIB-SEM.

### Focused Ion Beam Scanning Electron Microscopy (FIB-SEM)

#### Zebrafish

The resin block containing zebrafish eyes was prepared using a transmission electron microscopy (TEM)-adapted protocol provided in an earlier publication [17]. The serial section imaging was performed on a dual-beam FIB-SEM Helios 5 CX (Thermo Fisher Scientific Inc.) equipped with a Gallium liquid metal ion source. The resin blocks were mounted onto a standard SEM stub using a conductive epoxy resin glue (Ted Pella, #16014) and with silver resin paste to enhance sample conductivity (Ted Pella, #anchor16062). The stubs were carbon-coated using a Leica EM ACE600 sputter coater and loaded onto a multi-purpose holder in the SEM sample chamber. An overview low resolution back scattered electron image of the entire block was acquired with the Everhart-Thornley detector and the region of interest (ROI) was identified. The sample was next placed at the eucentric height and prepared for the Slice and Viewing workflow. Briefly, a protection layer of platinum (1–2 µm) was deposited on the top of the ROI using a gas injection system (GIS) with dimensions of 20 m (x) × 20 m (y) × 2 m (z). Subsequently, trenches were made in front of the ROI with dimensions of 60 m (x) × 30 m (y) × 15 m (z) and along each side (each 10 m wide) using the Focused Ion Beam. This process exposed the block face of the ROI. Once the site preparation was completed, the ROI was subjected to milling and imaging sequences using the Auto Slice & View 4.2 (ASV) software package. This software automated the serial sectioning and data collection process, enabling the acquisition of high-resolution 3D data. For each slice, 10 nm of the resin was removed at a 4 mm working distance using the FIB beam. SEM images were acquired using 2 kV accelerating voltage and beam current of 0.69 nA with the backscatter electrons (BSE) using the elstar in-lens detector (TLD) and the elstar in-column detector (ICD). SEM images were acquired at the sampling of 6 x 6 × 10 nm^3^ voxels with a dwell time of 10 µs; a total of 1750 slices were obtained. Imaged tissue dimensions are as follows: x=20.5 µm, y=15 µm, z=17.5 µm. The resulting stack of data was processed in Dragonfly 3D World (Comet Technologies Canada Inc., Version 2024.1) Briefly, we aligned individual images with a built-in “Stack Alignment” function, which corrected most of the major shifts that occurred during the acquisition. We then performed manual segmentation every 10-20^th^ section followed by an automatic Z interpolation with subsequent manual correction where necessary.

#### Mice

The FIB-SEM dataset used was previously described in Housset, et al. Briefly, mice were perfused intracardially with 10 mL of sodium cacodylate (Sigma-Aldrich, C0250) buffer 0.1 M, 0.1% CaCl2, pH 7.4, followed by 10 mL of 1% fixative solution (1% PFA; 1% (vol/vol) glutaraldehyde (Sigma-Aldrich, 354400) in sodium cacodylate buffer, and finally with 10 mL of a 2% fixative solution (2% PFA; 1% (vol/vol) glutaraldehyde in sodium cacodylate buffer). The dorsal pole was marked, and the eyes were enucleated and immersed overnight (ON) in 2% fixative solution + 10% sucrose at 4°C. The 1 mm^3^ of the dorsal retina was dissected and washed 3 times with sodium cacodylate buffer for 30 min, post-fixed in 0.1 M sodium cacodylate buffer containing 1% (wt/vol) Osmium tetroxide (OsO_4_) (Sigma-Aldrich, 201030) and 1.5% (wt/vol) potassium ferrocyanide (Sigma-Aldrich, 104984) for 2 h at 4 °C. After 3 washes of sodium cacodylate buffer for 30 min, the samples were immersed for 1 hour at 4°C in sodium cacodylate solution and 1% tannic acid (Sigma-Aldrich, 16201). Samples were rinsed 3 times in distilled H2O for 30 minutes and en bloc stained for one hour with 1% (wt/vol) aqueous uranyl acetate (Sigma-Aldrich, CDS021290) at 4 °C.

Samples were dehydrated in five successive steps of acetone and dH2O [30 to 90% (vol/vol)], each for 15 min at RT followed by 100% acetone (3 × 20 min). The samples were incubated with increasing concentrations [30 to 100% (vol/vol)] of low viscosity EPON 812 replacement (Mecalab Ltd, 3137N) over a period of 24 h, and then polymerized at 65 °C for 48 h. Ultrathin sections (70–100 nm) were en face cut from the resin blocks using a Leica Microsystems EM UC7 ultramicrotome (Leica Microsystems, Wetzlar, Germany) with a Diatome diamond knife (Diatome Limited, Nidau, Switzerland) and stained with 1% toluidine blue (Sigma-Aldrich, T3260) to ensure the quality of the preparation and locate the region of interest (ROI) before data collection by FIB-SEM.

The blocks were trimmed with a razor blade to expose the ROI, mounted on fixed 45° pre-tilt SEM stubs, and coated with a 4-nm layer of platinum using a Leica Microsystems EM ACE600 sputter coater (Leica Microsystems, Wetzlar, Germany) to enhance electrical conductivity. Milling of the serial sections and imaging of the block face after each Z-slice was carried out by the Helios Nanolab 660 DualBeam using Auto Slice & View G3 ver 1.2 software (Thermo Fisher Scientific). The sample block was first imaged to determine the orientation of the block face and ion and electron beams. A 2-m layer of platinum was deposited on the surface of the ROI to protect the resin volume from ion beam damage and correct for stage and/or specimen drift, i.e. orthogonal to the block face of the volume to be milled. Trenches on either side of the ROI were milled to minimize redeposition during the automated milling and imaging. Fiducials were generated for both the ion and electron beam imaging and used to dynamically correct for drift in the x- and y-directions during data collection by applying appropriate SEM beam shifts. Milling was performed at 30 kV with an ion beam current of 9.3 nA, stage tilt of 4°, and a working distance of 4 mm. At each step, a 23 nm slice of the block face was removed by the ion beam. Each newly milled block face was imaged with the through-the-lens-detector (TLD) for backscattered electrons at an accelerating voltage of 2 kV, beam current of 0.40 nA, stage tilt of 42°, and a working distance between 2.5 to 3.5 mm. The pixel resolution was 23.6 nm with a dwell time of 10 s per pixel. Pixel dimensions of the recorded images were 1536×1034. Nine-hundred and fourteen images were collected. Visualization and direct 3D volume rendering of the acquired datasets was done with Amira for Life Sciences software (Thermo Fisher Scientific).

Cone photoreceptors were distinguished from rod photoreceptors based on the difference in inner segment size and shape of the outer segment. Photoreceptor inner-segment membranes and Müller glial apical processes were manually segmented in successive z-planes using Imaris software (V10.2, Oxford Instruments). Missing in-between z planes were linearly interpolated.

### Genetic manipulation

#### CRISPR/Cas9 injections

All reagents for CRISPR/Cas9 injections were obtained from Integrated DNA Technologies, unless indicated otherwise. We ordered crRNAs targeting the third exons of *tbx2a*(TAACGATATGAAACCTGGGTTGG; GACAGCTATAAAATCGGTCTCGG) and *tbx2b* (TATCGTTGGCTCTCACAATATGG; CAAGGTATGTACCCATATTTTGG), that were previously designed and shown to successfully inhibit generation of UVS cones in F0 fish [26]. To prepare the gRNA:Cas9 complex, we followed a previously described protocol, with a few minor modifications [27]. Briefly, we diluted each ordered RNA to 100 M in IDT buffer and stored them at −20 °C. The day before the injections, we annealed crRNA with tracrRNA in a PCR machine with a program of 5 min at 95 °C, 0.1 °C/s cooling to 25 °C, 5 min at 25 °C followed by cooling to 4 °C and hold. On the morning of the injection, we diluted the crRNA:tracrRNA duplex with the IDT buffer to 25 M. Cas9 protein (stored at −20 °C) was also diluted to 25 M with RNAse-free phosphate-buffered saline (PBS), and then 2 L of the enzyme was mixed with the pre-annealed gRNAs (2 L of each duplex) and H2O to 10 L. The 5 M RNP complex was then incubated for 5 min at 37 °C and promptly brought to the injection room. Approximately 1 nL of mixture was injected into the yolk of *Tg(sws1:GFP)* embryos at one-cell stage using a Pneumatic PicoPump (PV820, WPI).

#### Generating *gfap*:GFP F0 fish

The Tol2 *gfap*:GFP construct was a gift from the laboratory of Dr. Ryan MacDonald. After transforming the bacteria (One Shot TOP10 chemically competent cells, Invitrogen), purifying the construct, and confirming the sequence, it was injected together with Tol2 mRNA into wild-type embryos, as described above. The embryos were selected for the experiments at 2 dpf under a fluorescent microscope based on GFP expression in the heart. The larvae were euthanized in the morning of days 3, 4, 5, 6, and 7.

#### *In vivo* subretinal electroporation of mouse RPE

*In vivo* subretinal electroporation of neonatal mice (P0) was done as previously described [28, 29] using 1 µl of plasmid DNA at 3 µg/ul. Mice pups were anesthetized on ice for 5 mins prior to the electroporation and allowed to recover in a heated chamber for 10 mins following electroporation. Immediately following subretinal injection, the treated eyes were subjected to electroporation with the positive electrode placed over the non-treated eye to drive the negatively charged plasmid DNA toward the RPE cells. A BTX electroporator ECM830 was used with the following settings: 80 volts, 50 ms pulse duration, 6 pulses, 0.95 sec pulse interval, 10 mm electrode gap.

#### Vectors

The plasmid driving expression of mNeonGreen-CAAX was derived from the Addgene vector pAAV-CAG-mNeonGreen (#99134) by insertion, in frame at the C terminus of the mNeonGreen coding sequence, of a nucleotide sequence encoding the classic Ras-derived C-terminal membrane-targeting tail (YGDGKKKKKKSK-TKCVIM), including the CAAX motif CVIM, using the BsrGI restriction site. The plasmid driving expression of mScarlet3-CAAX was derived from the Addgene vector pAAV-CAG-mTagBFP2-P2A-EGFP by replacing the mTagBFP2-P2A-EGFP coding sequence with the mScarlet3-CAAX coding sequence from the Addgene vector pLV-mScarlet3-CAAX-IRES-Puro (#222639), using the BamHI and EcoRI restriction sites. In both plasmids, transgene expression was driven by the CAG regulatory cassette, composed of the CMV enhancer, chicken beta-actin promoter, and rabbit beta-globin intron, enabling strong ubiquitous expression in mammalian cells.

### 0.1 Tissue processing for immunohistochemistry

#### Zebrafish

Zebrafish tissue preparation was performed as previously described [17]. For 3.5 mpf adults, instead of fixing euthanized fish whole in 4% paraformaldehyde (PFA), the eyes were extracted first, the cornea was cut, and the lens was removed. The eyes were then placed in 4% PFA and the regular protocol was followed. After three PBS washes and two sucrose cryopreservation steps, the fish/eyes were positioned in plastic molds, frozen in optimal cutting temperature compound, and stored at −80 °C (Tissue-Tek, Sakura Finetek). Blocks were cut on CM1900 cryostat (Leica), and 12 µm tissue sections were transferred onto Superfrost Plus slides (Fisherbrand). The slides were warmed and processed in a humidified chamber. The tissue was outlined with a lipid pen (Liquid Blocker, EMS) and rehydrated with PBS. Next, three permeabilization wash steps with PDT (0.1% Triton X-100 and 1% DMSO in PBS) were performed. For phosphorylated Ezrin, Radixin,and Moesin (pERM) stainings, protein-induced epitope retrieval using 0.05% Pronase (Millipore) in PBS was performed instead. Pronase solution was carefully pipetted onto the slides, which were incubated for 10 min at 37 °C and for another 10 min at room temperature, followed by three washes with PDT. Both regular and epitope-retrieved sections were further processed according to the previously published protocol [17]. Primary and secondary antibodies as well as fluorescent dyes can be found in Table S1.

#### Mice

Mice were sacrificed between post-natal days 40 and 50. All animals were maintained in a traditional 12-hour light-dark cycle. Light-adapted animals were sacrificed in the light and dark adapted animals were placed into a dark cabinet overnight and sacrificed in the dark. For both conditions, eyes were enucleated and fixed in 4% paraformaldehyde (PFA) (Electron Microscopy Sciences, 15710) for 1 hour at room temperature. For immunohistochemistry on retinal cross-sections, eyes were washed 3 times for 10 minutes each with PBS, incubated in 20% sucrose overnight, embedded in OCT (Tissue-Tek), frozen in liquid nitrogen, and stored at −80°C until sectioning at 18 µm (for retinal purposes) or 30 µm (for RPE purposes). For immunohistochemistry on retinal or RPE wholemounts, eyes were rinsed three times for 10 minutes each following fixation, and dissected to isolate the retina and RPE/sclera. Incisions were made to allow flattening of the retina (4 incisions) or RPE (8 incisions). Samples were permeabilized in blocking solution (0.3% Triton X-100 + 3% bovine serum albumin) overnight at room temperature. Primary and secondary antibody incubations were done overnight at room temperature in the blocking solution. Samples were washed in PBS between antibody incubations, and before mounting. After immunostaining, samples (cross-sections and wholemounts) were mounted flat in Mowiol (Millipore Sigma catalog #475904).

### Fluorescent imaging and processing

#### Zebrafish

Retinal images were obtained using a Zeiss LSM700 microscope with a 63x/1.4 NA oil objective and ZEN 2009 software. A Zeiss Elyra 7 Lattice SIM with a 63x/1.4 NA oil objective and ZEN 3.0 black were used to show pERM and glial GFP colocalization (Fig 1B). ImageJ (1.54p, [30]) was used to process fluorescent images. Multichannel plot profile in BAR plugin [31] was used to analyze the intensity of the RPE label at the OLM in Fig 2. For Fig 1A as well as S1C and D, the colocalization was obtained in ImageJ by setting threshold and creating a binary mask of both channels and then identifying the overlap using Image Calculator.

**Figure 1.**
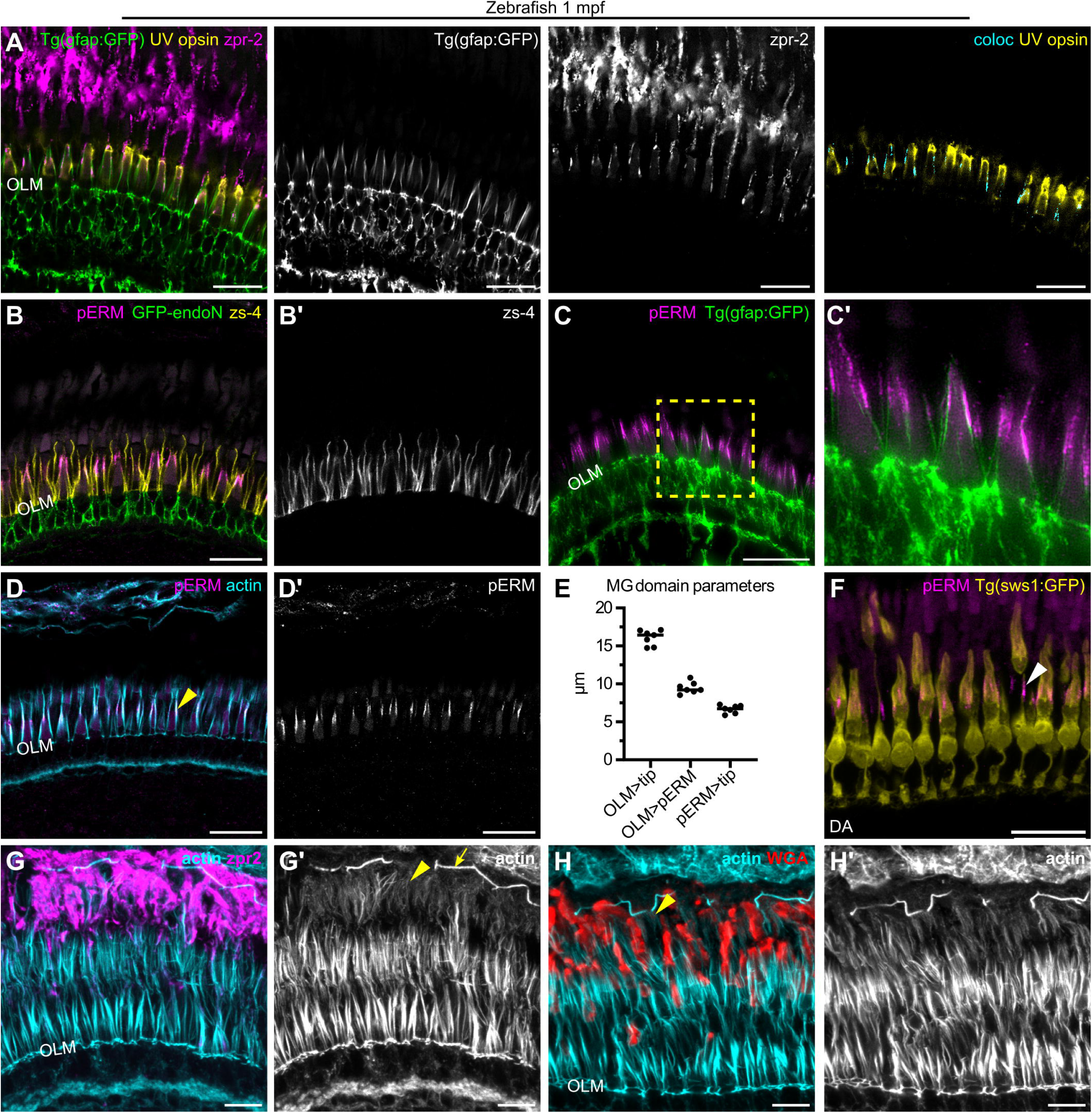
Characterizing apical processes of photoreceptor support cells. Images of the photoreceptor layer in immunostained cryosectioned 1 mpf zebrafish retina. (A) *Tg(gfap:GFP)* retina (GFP—green) labeled with zpr-2 antibody (magenta) to highlight glia–RPE colocalization (cyan) around UVS cones (anti-UV opsin, yellow); n=3. (B) Anti-pERM antibody (magenta), endoN-GFP (green), zs-4 (yellow), n=3. Zs-4 staining is also shown in a separate panel (B’). (C) *Tg(gfap:GFP)* retina (GFP—green) co-labeled with anti-pERM antibody (magenta), n=5. A zoomed in inset from the yellow box is also shown separately in (C’). (D) Colocalization of pERM (magenta) and actin stained with phalloidin (cyan) in thick bundles near the tip of the UVS outer segment (arrowhead), n=6. pERM staining is also shown in a separate panel (D’). (E) Graph highlighting the portion of the Müller glial processes labeled by pERM. Median is shown. (F) 1 mpf dark-adapted *Tg(sws1:GFP)* retina (GFP—yellow) stained with pERM (magenta). Arrowhead highlights a glial process not following a particularly elongated UVS cone. (G,H) 1 mpf crystal zebrafish retina stained with phalloidin (cyan); maximum intensity projection of a z-stack is shown, n=3. Brightness of the actin channel is increased to highlight diffuse filaments in proximal RPE processes. (G) RPE is visualized with zpr-2 (magenta); arrow in G’ outlines the actin at the tight junctions, whereas the arrowhead points at the actin in RPE processes. (H) Rod outer segment are stained with WGA (red); arrowhead shows actin localized to the RPE and not rod microvilli. Scale bars: 20 µm (A–D,F), 10 µm (G,H).

**Figure 2.**
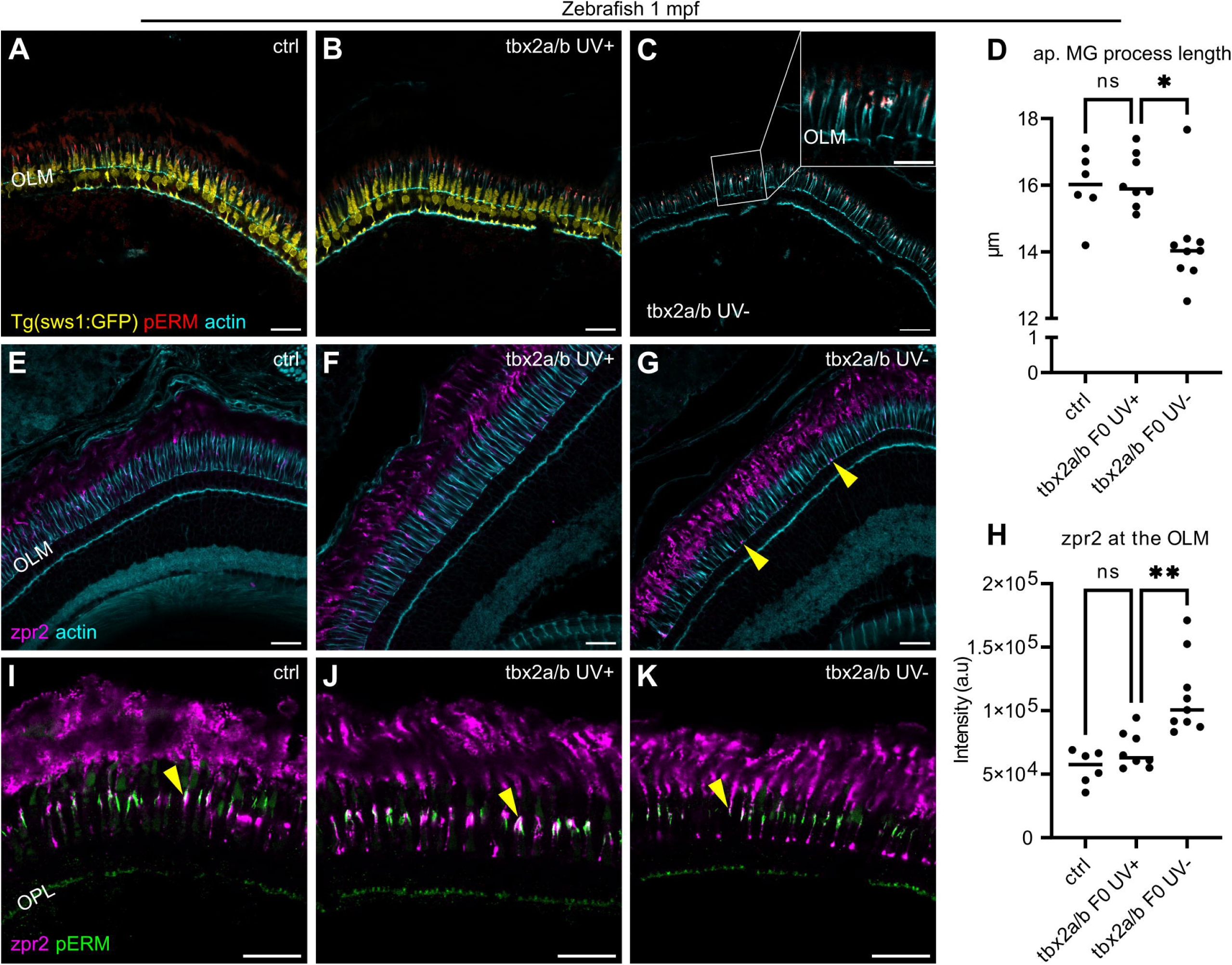
Apical Müller glial and RPE processes persist in 1 mpf zebrafish lacking UVS cones. (A–C) Images of *Tg(sws1:GFP)* zebrafish retinal sections (GFP—yellow) stained with phalloidin (cyan) and pERM (red); control (A), *tbx2a/b* UV+ (B), and *tbx2a/b* UV-(C). Inset in (C) shows a magnified view of the pERM staining. (D) Graph with length of apical Müller glial (MG) processes measured from the OLM to the tip of pERM signal and plotted together. (E–G) Control (E), *tbx2a/b* UV+ (F), and *tbx2a/b* UV-(G) retinal images labeled with actin (cyan) and zpr-2 (magenta). Arrowheads highlight enlarged distal tips of RPE processes in UV-retina. (H) Graph showing the intensity of the zpr-2 signal measured at the OLM. Statistics for both graphs: Welch’s ANOVA test, Dunnett’s T3 multiple comparisons test, median is shown. Number of eyes n=6 (control), n=8 (*tbx2a/b* UV+), n=9 (*tbx2a/b* UV-). (I–K) Control (I), *tbx2a/b* UV+ (J), and *tbx2a/b* UV-(K) retinas labeled with pERM (green) and zpr-2 (magenta). Arrowheads point at areas of overlap. Scale bars: 10 µm (inset), 20 µm.

#### Mice

Retinal cross-section images were acquired using the Zeiss LSM700 confocal microscope or Leica Stellaris confocal microscope at 40x magnification. Z-stacks and single section images were processed using ImageJ software. Retinal and RPE wholemounts were acquired using the Leica Stellaris confocal microscope at 96x magnification with Z-stacks taken at different intervals. Imaris Image Analysis Software was utilized for image processing and 3D reconstruction.

### Data Analysis

#### Zebrafish

GraphPad Prism software (10.5.0) was used to run statistical tests and to create graphs. Valid for all graphs: two-tailed p-value; ns, not significant (P>0.05), *P<0.05, **P<0.01, ***P<0.001, ****P<0.0001. In Fig 2 (FIB-SEM data), individual measurements are plotted. For Fig 1 and 5, each dot on the graph represents a fish. In Fig. 2, each dot represents an individual eye. When measuring zpr-2 signal at the OLM (Fig. 2), the analysis was blinded.

#### Mice

Quantifications are reported as mean ±SD and all figures were created using GraphPad Prism ver.9. Reported n values represent the number of biological replicates (independent animals). Animal numbers, cell counts and statistical methods are reported within the figure legends.

## Results

### Müller glial apical processes are molecularly partitioned and structurally stable in zebrafish

The zebrafish retina contains four subtypes of cones, typically labelled as red, green, blue, and UVS, in order from tallest to shortest (Figure S1B). We previously observed via confocal microscopy how Müller glia and the RPE protrude into the subretinal space and surround the UVS cone outer segments (Fig 1A) [17]. To better characterize the long apical processes extending from both cell types, we performed a series of immunostainings in the outer retina of 1 month post-fertilization (mpf) zebrafish.

The long apical glial processes above the OLM can be visualized with the *gfap*:GFP transgene or phalloidin staining of F-actin (Figure S1C) [17, 32, 33]. However, *gfap*:GFP requires access to a genetic line and phalloidin also labels actin in photoreceptors. Therefore, we sought to identify new labels and to better characterize process structure by examining a panel of alternative markers. We started with the common glial markers anti-glutamine synthetase (GS) and the zebrafish zrf-1 antibody, neither of which labelled Müller glia above the OLM (Figure S2A,B) [32, 34, 35]. The zs-4 antibody targeting the apical polarity marker Crb2a was previously shown to label apical Müller glia, which we confirmed (Figure S1D). Crb2a expression was present only in apical processes, above the OLM. However, zs-4 also labels Crb2a on photoreceptor inner segments [32], see Fig 1B and S1B. As we previously showed, the polysialic acid marker EndoN is specific for glia, localizing to the area between the OLM and slightly below the photoreceptor synaptic layer, but does not extend into the processes in zebrafish [36, 37]. In summary, the apical Müller glial region adjacent to photoreceptor nuclei is EndoN+, while the processes above the OLM are phalloidin+/zs-4+.

The actin-binding protein Ezrin was reported to localize to glial processes above the OLM in multiple species [38–40], and we confirmed phosphorylated Ezrin, Radixin, and Moesin (pERM) presence in the upper portion of the GFP-positive glial apical processes and absence in the apical domain of the RPE in the *Tg(gfap;GFP)* retinas (Fig 1C). When co-staining with phalloidin, we observed pERM signal colocalizing with the thickest actin bundles around the tip of the UVS outer segment (Fig 1D). Upon measurement, the pERM label was found to demarcate the upper 40% of the glial process above the OLM (Fig 1E).

To inspect the actin arrangement in the RPE cells, co-labelling with phalloidin and the RPE marker zpr-2 [41, 42] was performed on retinas from pigmentless *crystal* fish [25]. We observed actin filaments extending from the RPE cell bodies between rod outer segments (Fig 1G,H)). Still, the RPE phalloidin staining appeared much more diffuse and lower intensity than that observed in photoreceptors or glia. Of note, photoreceptors possess actin-based microvilli that extend alongside the outer segment, also known as calyceal processes. Due to the high actin expression in calyceal processes and their roots, it was not fully clear whether the RPE processes retained F-actin as they reached deeper into the subretinal space; however, we could not detect F-actin in their apical-most portions. Thus, unlike the prominent and organized F-actin in calyceal processes and Müller glia [17], we observed only thin and diffuse F-actin staining in the basal portion of the RPE extensions.

In teleost fish, including zebrafish, photoreceptors undergo shape changes in response to alternating light/dark conditions, as part of the so-called retinomotor movement mechanism. Cone inner segments elongate while rod inner segments shorten during dark adaptation, and the opposite occurs during light adaptation [43]. Notably, select UVS cones elongate dramatically in the dark, placing their outer segment almost fully above the outer segments of other UVS cones [17]. When tracking these particularly elongated UVS cones in the dark-adapted 1 mpf zebrafish retina, we did not observe Müller glial processes following the outer segment translocation (Fig 1F). Further, we specifically tested whether the actin core of glial processes remodels during dark adaptation to allow for extension alongside lengthening UVS cones. Eyes were analyzed from adult *Tg(hsp:act-myc)* zebrafish following a two-hour dark adaption post heat shock and timed to ensure Myc-tagged G-actin would be available during the period of retinomotor movements. As a control, the light-adapted group underwent heat shock but was maintained in lit conditions prior to fixation. Non-heat shock controls were also included for both dark- and light-adapted conditions. Consistent with our previous results, we observed the incorporation of newly expressed tagged actin into calyceal processes and inner segments in both light-adapted and dark-adapted retinae following heat shock, and minimal background staining in the non-heat shock controls (Figure S2D–G). In contrast, actin-Myc did not colocalize with the thick actin bundles of the glial processes surrounding UVS cones in either condition, suggesting that the Müller glia above the OLM remain unchanged during modifications of photoreceptor heights through the light cycle.

In summary, we demonstrated that the phalloidin+/zs-4+ long apical processes belonging to Müller glia can be subdivided into distal and proximal domains depending on the presence or absence of pERM immunoreactivity, respectively. Furthermore, our data suggest that the glial processes are stable and feature a robust core of aligned actin bundles, whereas the RPE processes contain diffuse actin only proximally.

### Müller glia and the RPE remain in contact in the absence of UVS cones

The glia–RPE interaction in zebrafish is concentrated around UVS cones, which led us to ask whether they are necessary to facilitate contact between the two support cell types. To address this question, we examined zebrafish lacking UVS cones (Fig 1A). Earlier work highlighted two paralogous T-box transcription factors (Tbx), *tbx2a* and *tbx2b*, as being essential for differentiation of short single UVS cones [26]. We co-injected the previously published gRNAs together with Cas9 enzyme into *Tg(sws1:GFP)* zebrafish embryos. We then analyzed 1 mpf F0 crispants, comparing injected fish lacking UVS cones to injected fish with cones remaining and uninjected controls.

In five out of nine analyzed injected fish, the UVS cones were almost entirely absent (*tbx2a/b* UV-), with only a few morphologically aberrant GFP-expressing cones remaining (Figure S2H). To check for random silencing of the *sws1*:GFP transgene, we co-labeled the tissue with an anti-UV opsin antibody, confirming the absence of normal UVS cones. Interestingly, in fish lacking UVS cones, UV opsin was detected within taller non-UVS cones, a staining pattern not present in either control fish or *tbx2a/b* crispants with UVS cones preserved (*tbx2a/b* UV+; Figure S2J–L).

The long apical processes of Müller glia and RPE cells persisted and were closely aligned in the retinas lacking UVS cones (Fig 2). We measured the length of the glial apical processes from the OLM to the tip, labeled by pERM (Fig 1, Fig 2A–D). While Müller glial processes were statistically significantly shorter in *tbx2a/b* UV-fish, the reduction only amounted to around 11% of the length in the two control groups.

In addition, we observed that the RPE processes in the *tbx2a/b* UV-retinas appeared enlarged at their distal tips, near the OLM (Fig 2E–G). We compared the size of the RPE processes immediately apical to the OLM in *tbx2a/b* UV- and *tbx2a/b* UV+ by measuring the intensity of zpr-2 in a defined region around the OLM, Figure S3A–D. When plotting the intensity of the peaks, we observed a significant increase in zpr-2 signal in *tbx2a/b* UV-retinas, compared to the two control groups (Fig 2H), suggesting an expansion of the tips of the RPE apical processes in the absence of UVS cones. In the two control groups, the RPE (together with glial processes) instead formed a cap over the UVS outer segment tip (Fig 2I–K and Figure S2I).

Altogether, we discovered that UVS cones are dispensable for the extension and alignment of RPE and Müller glial apical processes in the subretinal space.

### CD44-positive Müller glial apical processes extend through the subretinal space and associate with cones in mice

While we demonstrated glia–RPE alignment surrounding UVS cones in fish, it is unclear whether this relationship is conserved in other species, in particular in the rod-dominated mouse retina. To investigate whether long glial processes above the OLM are present in mice, we examined adult mouse retinal cross-sections and flatmounts by immunofluorescence using general cytoskeletal probes together with glia-specific markers. Unlike in zebrafish, where phalloidin and pERM prominently label Müller glia but not RPE apical processes, both apical compartments were detected by these probes in the adult mouse retina (Fig 3A). In both Müller glia and RPE cells, phalloidin and pERM showed distinct but comparable distributions. In Müller glia, phalloidin extended broadly from the junctional OLM region to the distal tips of apical processes, whereas pERM was enriched more proximally, immediately above the junctions, and progressively weakened distally (Fig 3A, Figure S4). To establish the cellular identity of these phalloidin-positive structures, we next used CD44, a transmembrane glycoprotein and established marker of the Müller glial apical surface [44]. CD44 labeled thin apical processes emerging above the OLM and colocalized with the phalloidin-positive processes extending into the basal subretinal space, confirming their Müller glial origin (Fig 3B).

**Figure 3.**
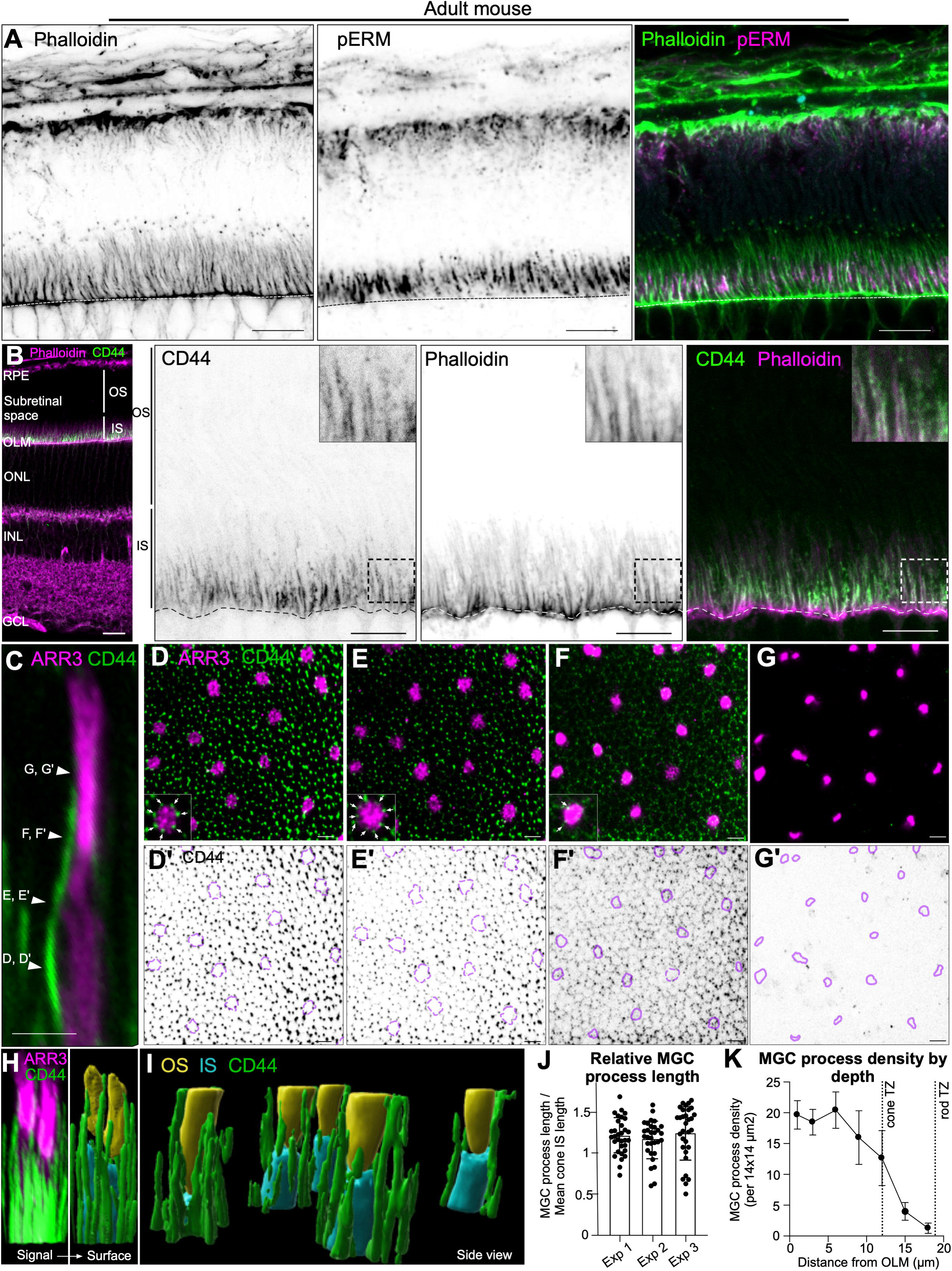
CD44 labels Müller glial apical processes associated with cones in the mouse subretinal space. (A) Immunostainings of an adult mouse retina cross section through the outer retina. Phalloidin and phospho-ERM (pERM) label apical processes of both Müller glia (MG) and RPE cells. The dashed line indicates the outer limiting membrane (OLM). Scale bar: 10 µm. (B) Immunostainings of an adult mouse retina cross-section showing colocalization of the MG apical process marker, CD44 (green), with the F-actin marker phalloidin (magenta), in its basal region at low magnification (left) and high magnification (right). The dashed line indicates the OLM. RPE: retinal pigment epithelium, OLM: outer limiting membrane, OS: outer segment, IS: inner segment, ONL: outer nuclear layer, INL: inner nuclear layer, GCL: ganglion cell layer. Scale bars: 20 µm (left panel), 10 µm (right panels). (C) High magnification of a single cone (ARR3, magenta) and an associated MG process (CD44, green). Arrowheads indicate placement of optical sections for panels D,D’–G,G’. Scale bar: 2 µm. (D–G) En face images of whole-mount adult mouse retina immunostained for ARR3 and MG apical processes (CD44), acquired at increasing distances from the OLM. Arrows in the magnified inset highlight MG processes in direct contact with cones, located within ARR3 indentations at the inner segment (IS) level. Scale bar: 3 µm. (D’–G’) Corresponding images to panels D–G with the CD44 signal inverted. Cone IS and outer segment are outlined with dashed and solid lines, respectively. (H) Representative 3D fluorescence dataset showing CD44 (green) and ARR3 (magenta) signals (left), together with the corresponding surface-rendered reconstruction used to segment cone inner segments (IS; blue), outer segments (OS; yellow), and Müller glial processes (green) (right). (I) Independent 3D surface reconstruction focused on the cone transition zone, showing Müller glial processes (green) extending along the cone IS (blue) and reaching the base of the cone OS (yellow). (J) Quantification of Müller glial process length relative to mean cone inner segment (IS) length in the corresponding retinal region. Each dot represents a single MG process normalized to the mean cone IS length of the same retina. Each distribution represents one retina, for a total of three retinas analyzed. (K) Quantification of MG process density as a function of distance from the OLM (n=3 retinas). Dashed lines indicate the average depth of the photoreceptor IS-to-OS transition zone (TZ) based on photoreceptor morphology established in the literature, corresponding to the average inner segment length of 12 m for cones and 19 m for rods [45, 46].

Retinal cross-sections showed that CD44-positive Müller glial processes densely populate the subretinal space. These filamentous glial processes extend from the OLM between both rod and cone inner segments, without obvious differences in density, length, or overall organization (Fig 3C,C’). En face optical stacks further revealed that CD44-positive Müller glial processes form a dense basal network between cone inner segments, where they occupy stereotyped ARR3-negative indentations along the cone contour and remain associated with cones over increasing distances from the OLM (Fig 3D–G). Inverted renderings of the same planes emphasized the weaker distal CD44 signal as a function of distance from the OLM (Fig 3D’–G’). 3D reconstructions of the CD44 and ARR3 signals, together with surface rendering of cone inner segments and outer segments, confirmed that these Müller glial processes run alongside cone inner segments, and that a subset extend distally along cone outer segments (Fig 3H,I, and Video S1). Consistent with this finding, quantification showed an average apparent process length ratio to cone inner segment average length of approximately 1.2 (Fig 3J), while density measurements from the OLM demonstrated a progressive decline in the number of labeled structures with increasing depth, with many processes ending near the cone transition zone, some reaching the cone outer segment base, and few persisting to the mid-outer segment (Fig 3F,G,K).

Altogether, these data reveal a widespread network of long apical glial processes in the mouse subretinal space, with a subset reaching cone outer segments.

### Long RPE apical processes associate with cone photoreceptors and interact with Müller glia in mice

In zebrafish, our data showed that Müller glial and RPE apical processes are closely associated within the subretinal space, where they ensheath UVS cone outer segments. In the mouse retina, pERM and phalloidin staining similarly suggested that RPE-derived apical structures might interact with cones, as pERM-positive processes were observed running alongside cone outer segments labeled by the cone extracellular marker peanut agglutinin lectin (PNA) (Fig 4A). However, since no marker reliably and specifically labels the finest RPE apical membranes in the mouse retina, this potential RPE–cone interaction could not be definitively resolved by conventional immunostaining. We therefore used *in vivo* electroporation at post-natal day 0 (P0) to drive sparse expression of a membrane-targeted fluorescent protein (mNeonGreen-CAAX or mScarlet3-CAAX) in RPE cells [28, 29], enabling bright labeling of individual RPE cells and their apical and lateral processes (Fig 4B, Figure S5). Characterization of electroporated RPE cells confirmed the typical epithelial cell morphology and maintenance of phalloidin-positive F-actin tight junction structure, indicating that RPE electroporation does not alter endogenous cell morphology or structure (Figure S5).

**Figure 4.**
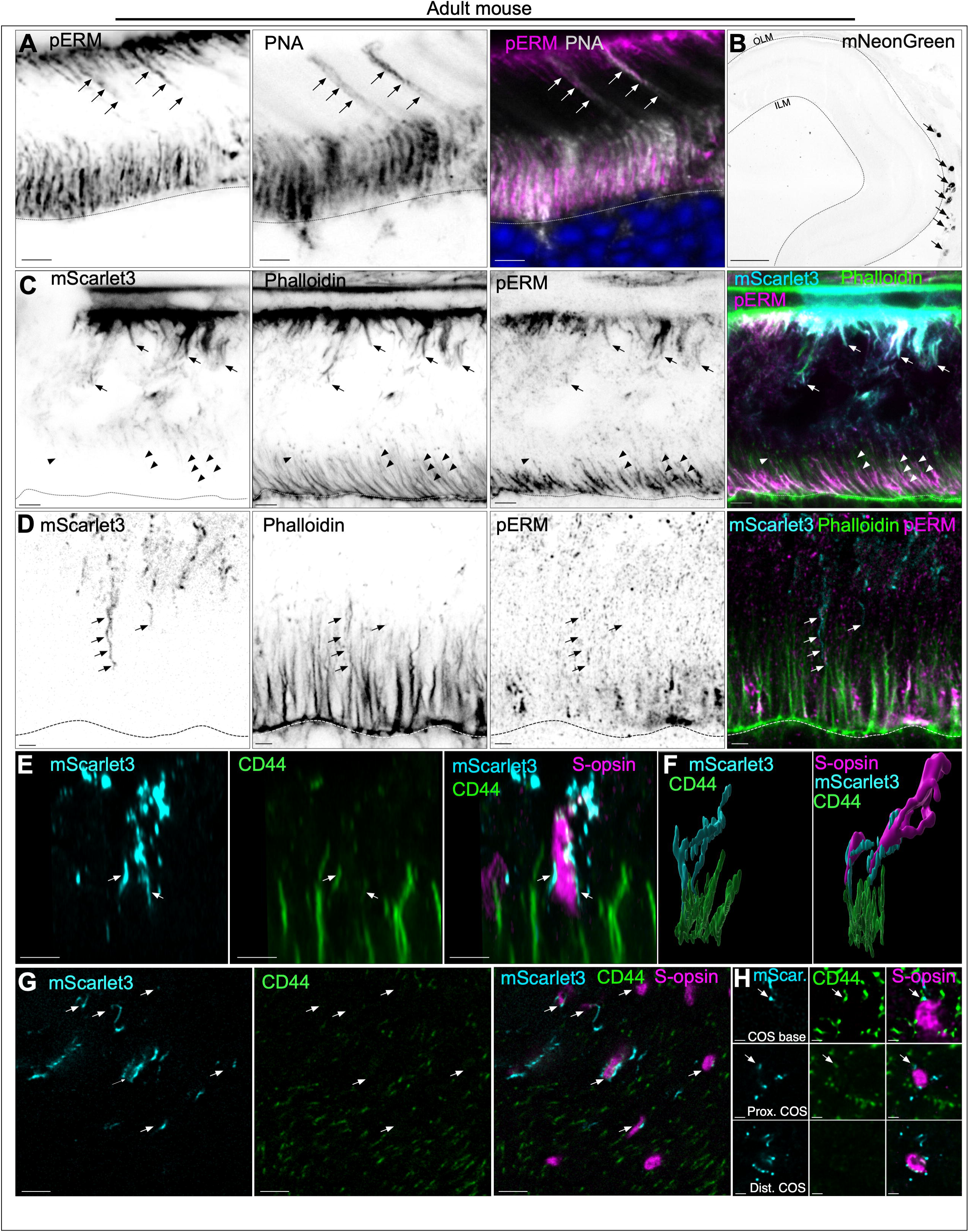
RPE apical processes extend into the basal mouse subretinal space and contact Müller glial apical processes around cones. (A) Immunostaining of an adult mouse retinal cross-section through the outer retina. pERM-positive processes extend alongside the outer segments (OS) of PNA-positive cones (arrows). The signal was inverted to enhance weak fluorescence. Scale bar: 5 µm. (B) Low-magnification mNeonGreen epifluorescence image of an adult mouse retina following P0 electroporation to sparsely label RPE cells with membrane-bound mNeonGreen-CAAX, revealing RPE cell morphology. The dashed lines indicate the OLM and inner limiting membrane (ILM). Scale bar: 200 µm. (C) Epifluorescence and immunolabeled images of phalloidin-stained adult mouse retinal cross-sections following P0 electroporation to sparsely label RPE cells with membrane-bound mScarlet3-CAAX (hereafter, mScarlet3-RPE). Both proximal (arrows) and distal processes (arrowheads) were labeled by mScarlet3-RPE. Dashed lines indicate the OLM. Scale bar: 4 µm. (D) Higher-magnification epifluorescence and immunolabeled images of phalloidin-stained adult mouse retinal cross-sections in the subretinal region. Arrows indicate mScarlet3-RPE apical processes extending into the basal subretinal region populated by phalloidin-positive Müller glia (MG) apical processes. Dashed lines indicate the OLM. Scale bar: 2 µm. (E) Epifluorescence and immunolabeled images of adult mouse retinal cross-sections in the subretinal region. mScarlet3-RPE processes (cyan) ensheath the OS of an S-opsin-positive cone (magenta) and contact CD44-positive MG apical processes (green). Arrows indicate regions of overlap between mScarlet3-RPE processes and MG processes surrounding a cone. Scale bar: 4 µm. (F) 3D reconstruction of mScarlet3-RPE distal processes (cyan) and CD44-positive MG apical processes (green), showing points of contact between the two surfaces (left). The same structures are shown together on the right with a 3D reconstruction of the S-opsin-positive cone OS (magenta), indicating that this RPE/MG interaction occurred at the position of a cone. (G) Epifluorescence and immunolabeled en face images of an adult mouse retina acquired in the plane of cone OSs. mScarlet3-RPE processes (cyan) were observed contacting S-opsin-positive cone OSs (magenta), independently of CD44-positive MG processes (green). All S-opsin-positive cones in the electroporated patch region were in contact with mScarlet3-RPE processes (arrows). Scale bar: 4 µm. (H) Epifluorescence and immunolabeled en face images of adult mouse retina acquired at three different planes along the cone OS axis (OS base, proximal OS, and distal OS). At the base of the S-opsin-positive cone OS (magenta), mScarlet3-RPE processes (cyan) were in close proximity to, and ran alongside, the distal tips of CD44-positive MG apical processes (green, arrow). These interactions were no longer observed in the distal cone OS region, where only the cone-RPE interaction remained. Scale bar: 1 µm.

Sparse RPE cell membrane labeling confirmed that the phalloidin- and pERM-positive apical structures arise from RPE cell bodies (Fig 4C). As in Müller glia, the proximal shafts of these RPE extensions were strongly labeled by phalloidin and pERM, whereas their most distal portions displayed much weaker signal, indicating that the actin-rich, pERM-positive domain is concentrated proximally, while the finest distal membranes are best revealed by membrane-bound fluorescent reporters (Fig 4C,D, Figure S5). This approach further uncovered a strikingly non-uniform organization of RPE apical processes in the mouse subretinal space which, importantly, remains unchanged in dark-adapted mice (Figure S6). In contrast to the widespread and relatively homogeneous glial network, RPE cells extended sparse, long apical processes that, in some cases, contacted the OLM (Figure S5H). These longer processes were preferentially associated with cones rather than spread evenly across the retinal plane, and contacted CD44-positive Müller glial apical processes (Fig 4E). 3D reconstruction confirmed direct points of apposition between distal RPE and Müller glia in the immediate vicinity of a cone outer segment (Fig 4F). En face imaging at the level of cone outer segments further showed that distal RPE processes are enriched in the retinal plane where cones are positioned (Fig 4G). Still, not every labeled RPE long process was associated with the position of a cone (Fig 4G). Furthermore, optical sectioning through successive planes of the cone outer segment axis showed that, at the outer segment base, RPE processes run alongside the distal tips of CD44-positive Müller glial apical processes. At the proximal cone outer segment, this relationship is maintained, but since glial processes do not extend longer, only RPE processes associate with the distal cone outer segment (Fig 4H).

Altogether, these data identify novel regions of interaction between Müller glia and the RPE at the site of cone locations in the mouse subretinal space. Unlike the broadly distributed glial network, long RPE apical extensions are spatially biased toward cone locations, where they contribute to a specialized tripartite interaction between cones, Müller glia, and the RPE. This cone-centered arrangement echoes the structural organization observed in fish. However, the mouse architecture appears less elaborate, with only a few glial processes reaching the cone outer segment base while multiple RPE apical processes surround the cone outer segment and occasionally contact those distal Müller glial extensions (Fig 4E–H, Video S2).

### Interaction between Müller glia and RPE in developing zebrafish

Visual function begins early in zebrafish development, with the first photoreceptor light responses detectable by 3 days post-fertilization (dpf) [47]. To understand when Müller glial long apical processes emerge and establish contact with RPE apical processes, we imaged ocular sections of larvae mosaically expressing GFP in Müller glia under the *gfap* promoter (Fig 5A–E, Figure S2C).

**Figure 5.**
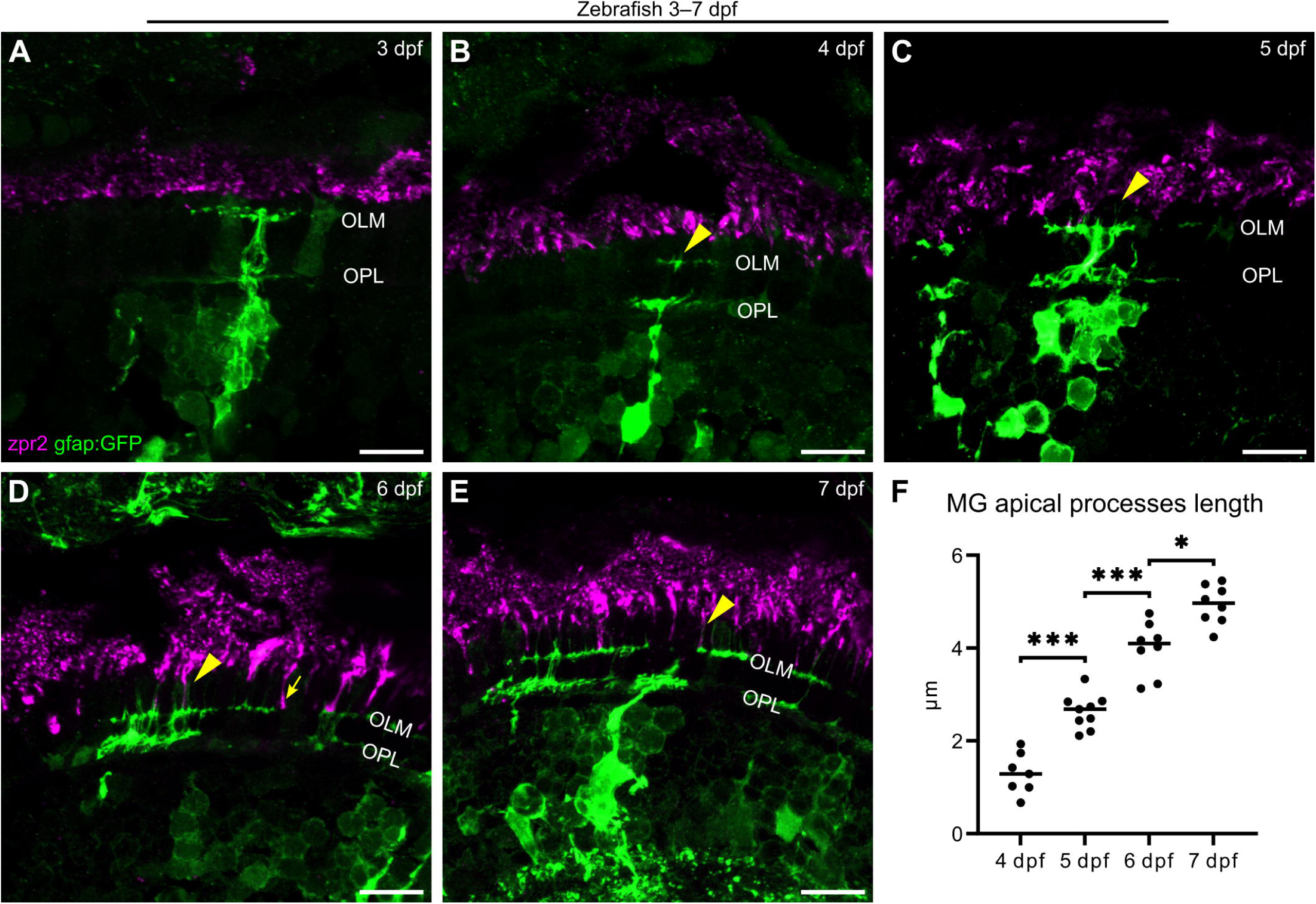
The establishment of contact between Müller glial and RPE apical processes in zebrafish. (A–E) Confocal images of larval retina with mosaic *gfap*:GFP expression (GFP—green, zpr-2—magenta). Progression of the interface between Müller glia (MG) and the RPE in the developing retina; maximum intensity projection of a z-stack is shown. Arrowheads point at MG processes above the OLM. Arrow in (E) highlights RPE processes descending to the OLM. Scale bars: 10 µm. (F) Graph showing the growth of MG apical processes during development. Welch’s ANOVA test, Dunnett’s T3 multiple comparisons test, median is shown. Number of fish: n=5 (3 dpf), n=7 (4 dpf), n=9 (5 dpf), n=8 (6, 7 dpf).

Previously, no Müller glial presence was observed above the OLM in zebrafish larvae at 3 dpf (when the outer segment is already formed); thus, this timepoint was chosen for the earliest stage of our analysis [17]. The progression of the apical glia–RPE interface in the developing retina is demonstrated in Fig 5A–E. In accordance with our earlier data, no apical glial processes were detected at 3 dpf (Fig 5A); however, at 4 dpf short Müller glial projections began to grow from the OLM in the direction of the RPE (Fig 5B,F). At 5 dpf, the projections became slightly longer, and RPE cells were now creating extensions towards the OLM (Fig 5C). Here, we also noticed RPE protrusions overlapping with Müller glia for the first time, although in an inconsistent manner. In 6 dpf larvae, we observed elongated apical RPE processes reaching the OLM (Fig 5D, arrow) and running alongside long thin Müller glial processes. By 7 dpf, the Müller glial processes elongated further, and the glia–RPE contacts became more numerous and extensive (Fig 5E).

Taken together, apical processes of both Müller glia and the RPE extend and begin to interact early during development, although subsequent to photoreceptor outer segment formation [48].

### Müller glial apical processes emerge before RPE apical processes in mice

Next, we investigated when Müller glial and RPE apical processes emerge and progressively elaborate their interactions during postnatal cone maturation in the mouse retina. Retinas from eyes electroporated at P0 with membrane-bound fluorescent reporters were collected at P7, P13, and P15, corresponding to pre-, peri-, and post-eye opening, respectively, and analyzed in cross-section and 3D surface reconstructions. The contralateral, non-electroporated eye was collected simultaneously to serve as a control for immunostaining (Figure S4). At P7, CD44-positive Müller glial apical processes were already present, extending above the OLM and spanning the full thickness of the still narrow subretinal space, where they terminated against the pERM-positive apical surface of the RPE (Fig 6A,B). At this stage, however, the RPE apical domain remained flat and had not yet elaborated obvious processes into the subretinal space (Fig 6A,B). Strikingly, although the RPE apical domain was already pERM-positive, the glial apical processes remained pERM-negative, indicating that these early Müller glial extensions form before acquiring the apical molecular specialization seen in the adult (Fig 6A). Cones are still immature at P7, with rudimentary outer segments and S-opsin distributed broadly along the cell membrane. CD44-positive Müller glial processes were the main structures contacting them, in some cases extending along the cone inner segment, with no evident cone-associated RPE processes at this stage (Fig 6C,D).

**Figure 6.**
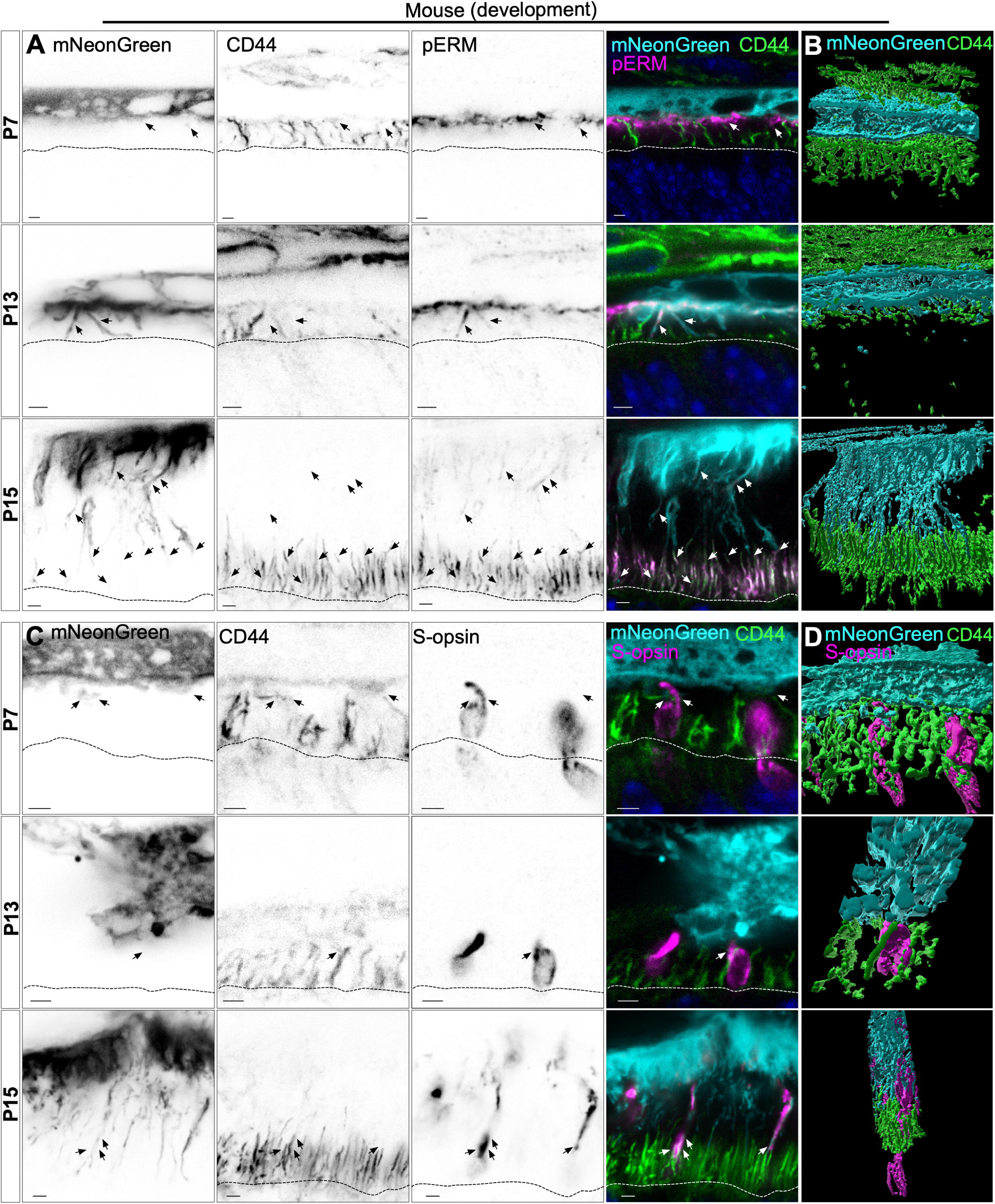
Müller glial apical processes emerge before RPE apical processes during cone maturation. (A) Epifluorescence and immunolabeled images of mouse retinal cross-sections following P0 RPE electroporation sacrificed at the indicated stages. Arrows indicate apical RPE processes. Immunostaining for pERM (magenta) highlights the actin dynamics of apical RPE processes (mNeonGreen, cyan) and Müller glial (MG) apical processes (CD44, green) during post-natal development. Dashed lines indicate OLM. Scale bar: 2 µm. (B) 3D surface reconstructions of CD44-positive MG processes (green) and mNeonGreen epifluo-rescence from mNeonGreen-RPE cells (cyan) at the indicated stages, highlighting the morphogenetic events shown in panel A. (C) Epifluorescence and immunolabeled images of mouse retinal cross-sections following P0 RPE electroporation and sacrificed at the indicated stages. Immunostaining for cones (S-opsin, magenta) allows to follow its interaction with RPE (mNeonGreen, cyan) and MG apical processes (CD44, green) during post-natal development. Arrows indicate interaction between MG and RPE processes at the vicinity of a cone. Arrowheads indicate cone-specific RPE apical processes. Dashed lines indicate OLM.Scale bar: 2 µm. (D) 3D surface reconstructions of CD44-positive MG processes (green), mNeonGreen epifluorescence (cyan) and cone S-opsin (magenta), at the indicated stages, highlighting the morphogenetic events shown in panel C.

At P13, both cellular compartments had elaborated apical processes that still remained immature. Electroporated RPE cells now had elaborated apical processes spanning the thickness of the subretinal space, and these emerging RPE extensions were pERM-positive, unlike the still pERM-negative CD44-positive Müller glial processes (Fig 6A,B). Occasional contacts between the two cell populations could already be observed, but cone organization at this stage remained broadly similar to P7, with rudimentary outer segments and glial processes remaining the main cone-contacting structure in the subretinal space (Fig 6C,D). Thus, the main developmental transition at P13 is the appearance of RPE apical processes, but not yet the establishment of a mature cone-RPE interface.

At P15, the subretinal space had markedly thickened and the mature organization began to emerge. CD44-positive Müller glial processes had now acquired pERM immunoreactivity, residing in the basal region of the subretinal space, while long RPE apical processes extended distally and contacted the tips of apical Müller glia (Fig 6A,B). At this stage, long mNeonGreen-RPE processes ran specifically along S-opsin-positive cone outer segments, whereas glial apical processes remained positioned more basally and contacted those RPE extensions at their tips, establishing the mature tripartite interface between cone, Müller glia, and RPE (Fig 6C,D). In addition, S-opsin-positive fragments could be detected along the upper tracks of RPE processes (Fig 6C), raising the possibility that cones and the RPE exchange biological material coinciding with the time of eye opening and the beginning of visual function. Together, these observations indicate that apical compartments of Müller glia and the RPE mature sequentially and form asynchronously during postnatal mouse development.

### FIB-SEM imaging reveals glia–RPE cage around UVS cones in zebrafish

To resolve the 3D organization of Müller glial and RPE apical processes around UVS cones, we performed FIB-SEM acquisition on juvenile zebrafish outer retina (Figure S7). The imaging captured several UVS cones from the nucleus or proximal inner segment to the distal outer segment. A central UVS cone was manually reconstructed, with the following classes traced: 1) inner segment including its continuation in form of calyceal processes and 2) outer segment and accessory outer segment, which together provided landmarks for mapping adjacent 3) Müller glial and 4) RPE processes. Müller glial processes were identified by the continuity with the distinct glial presence at the OLM, whereas structures we attributed to the RPE class contained electron-dense pigment granules. The resulting volume reconstruction is shown in Fig 7D–I and Video S3, while quantification of the glial and inner segment portions of the cage is presented in Fig 7C, J–L. Additionally, individual annotated images from different planes can be found in Figure S8A–G while animation of the whole stack is presented in Video S4.

**Figure 7.**
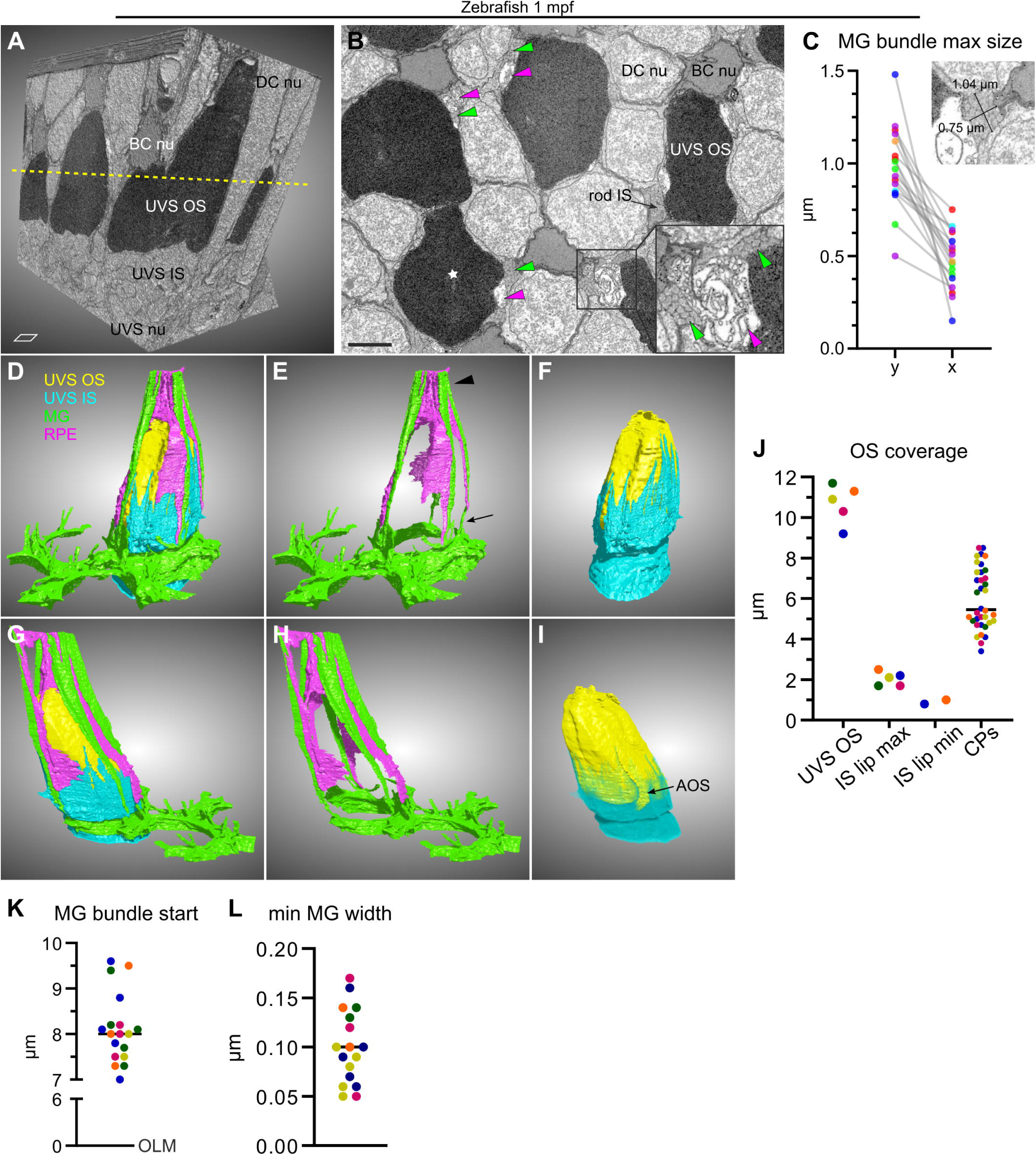
Müller glia, RPE, and calyceal processes form a cage around UVS outer segment in zebrafish. (A) A stack of all FIB-SEM sections shown cropped and in 3D view. The region captured ranges from the upper part of the UVS cone nuclei (nu) and inner segments (IS) to above the tip of the outer segments (OS). DC—double cone, BC—blue cone. A dashed line shows the approximate position of the plane shown in (B). Scale square side: 2 m. (B) A single section with photoreceptor types annotated. Green and magenta arrowheads point at several glial and RPE processes, respectively. Star signifies the UVS cone that was chosen for segmentation. Scale bar: 2 m. (C) Two dimensions of a MG process measured at its thickest point with an example. The longest axis was measured as y, with x perpendicular to y. (D-I) 3D reconstruction of the 4 segmented classes: UVS inner segment (IS)—cyan, UVS outer segment (OS)—yellow, Müller glia (MG)—green, and RPE—magenta. View from 2 sides is shown (D–F vs G–I). (E) Note the difference in diameter of the MG processes close to the OLM (arrow) and near the UVS OS tip (arrowhead). In (I), the IS opacity is reduced to highlight the accessory OS (AOS). (J) Length of the inner segment (IS) lip and calyceal processes (CPs) vs outer segment (OS) length. Lip min refers to a continuous IS sheet, without any gaps, and lip max represents an IS sheet with some gaps but which has not yet divided into individual CPs. Note that it was not feasible to reliably measure lip min in all UVS cones in the dataset. (K) The distance from the OLM where widening Müller glia (MG) processes begin to adopt a bundled organization. (L) Minimal width of MG processes, measured near the OLM. Within individual graphs, each colour represents measurements from a single cell. Medians are shown where appropriate.

Upon inspection of the reconstruction, we observed that the inner segment transition into calyceal processes did not occur at the inner segment–outer segment junction, as typically assumed. On the contrary, the first 1–2 m of the outer segment was almost fully covered by a continuous sheet of extended inner segment (Fig 7F,I,J). Furthermore, the length of the calyceal processes was not consistent, with the shortest ones measuring 3–4 µm and the longest ones at 8 µm reaching almost the entire outer segment height. Notably, the longest calyceal processes were located on the opposite side of the outer segment relative to the connecting cilium and accessory outer segment.

Müller glial and RPE processes were closely aligned and together encircled the UVS cone outer segment as 4–5 bundles positioned outside of the inner segment/calyceal processes ring (Fig 7 D,E,G,H). The glial processes mostly originated from a single cell and featured a remarkable transition in morphology as they extended from the OLM towards the tip of the UVS outer segment. Initially, Müller glial processes were very narrow, 0.1 m in diameter (Fig 7L), but expanded towards the tip of the outer segment (8 m from OLM) up to 1 m across and adopted a highly ordered ultrastructure with an organized bundled appearance, resembling multiple crayons tied together (Fig 7B,C,E,H,K, Figure S8A–G). Interestingly, we also observed rare short glial processes reaching only around the inner segment–outer segment junction. The RPE processes exhibited a less structured organization, were filled with membranous lamellae indicative of endoplasmic reticulum (ER), and harbored a few small pigment granules around the UVS outer segment tip (Figure S8G). To confirm ER presence in the apical RPE, we labelled zebrafish retina with a known ER marker, Calreticulin, which colocalized with zpr-2 around the UV cone outer segment tip (Figure S8H).

In summary, apical processes of the two cell types supporting photoreceptors—Müller glia and RPE— surround a UVS cone outer segment in a cage-like arrangement. Furthermore, the processes are closely aligned both alongside and above the outer segment, suggesting a more direct interaction between the support cells than previously appreciated.

### FIB-SEM imaging reveals Müller glial and RPE contacts around cones in mice

To further characterize the ultrastructural relationship between glia, RPE, and cones in mice discovered in fluorescent images, we analyzed a FIB-SEM dataset acquired from the outer retina of an adult mouse, spanning the region from the OLM to approximately the mid-length of rod outer segments [49]. The acquired volume contained eight cones together with neighbouring rods, RPE, and glial elements within the subretinal space. Briefly, a cone-centered subvolume was cropped from the dataset, and we reconstructed a central cone with its inner segment, outer segment, connecting cilium, and axoneme, as well as associated Müller glial and putative RPE processes. In parallel, the larger acquired volume was reconstructed for all classes except for RPE processes, whose fine morphology could not be resolved with sufficient confidence throughout the full dataset. The imaging layout of the full acquired volume is shown in Figure S9, Figure S10A,B, and Video S5 whereas the cone-centered subvolume used for fine-structure reconstruction is shown in Fig 8A and Video S6.

**Figure 8.**
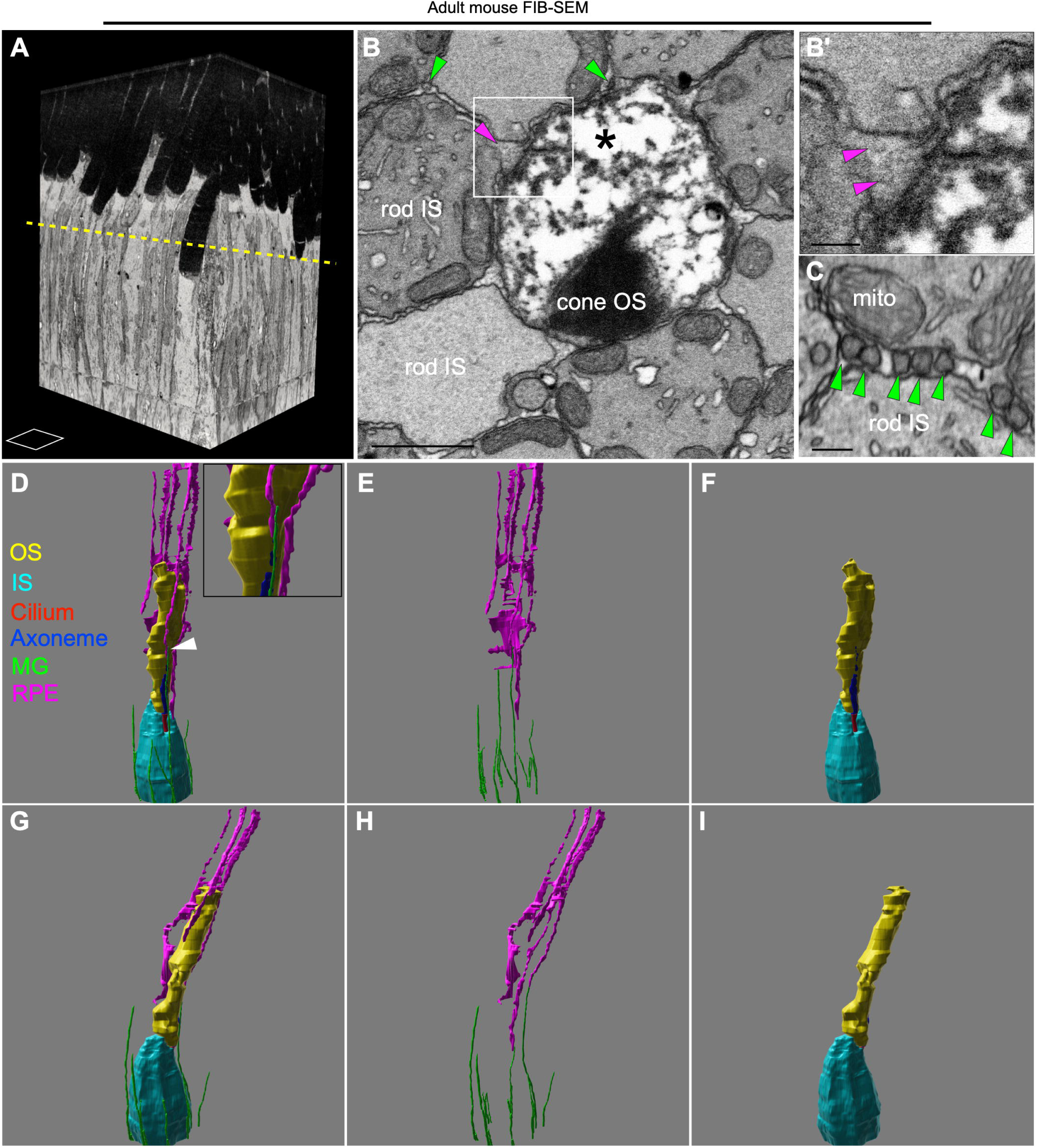
Müller glial and RPE apical processes contact cones in the mouse subretinal space. (A) A stack of all FIB-SEM sections shown as a cropped volume. The imaged area ranges from cone inner segment base to rod outer segment mid-length. A dashed line indicates the approximate position of the plane shown in (B). Scale square size: 4 µm^2^. (B) Single en face section with annotated photoreceptor types. Green and magenta arrowheads indicate several Müller glia (MG) and RPE processes, respectively. Scale bar: 1 µm. (B’) Enlarged view of the boxed region in (B). Scale bar: 200 nm. (C) High magnification acquisition at the level of inner segment base, showing the homogeneous round morphology of MG processes. Scale bar: 200 nm. (D–I) 3D reconstruction of the 6 segmented classes: cone inner segment (IS, cyan), outer segment (OS, yellow), connecting cilium (red) and axoneme (blue), MG (green) and RPE (magenta) apical processes. Two viewing angles are shown (D–F versus G–I). Arrowhead in (D) points to the interaction between the RPE and MG.

Glial processes were defined by their continuity with the Müller glial apical endfeet at the OLM, where vesicular structures with a whorled multilamellar appearance were also observed adjacent the base of the processes (Figure S10D). By contrast, structures assigned to the RPE class were identified more cautiously based on their position at the apical side of the subretinal space and on morphological features reminiscent of zebrafish RPE processes, including optically lighter vesicular profiles (Fig 7B; Fig 8B,B’,D–I; Figure S10F). The resulting 3D reconstruction is shown in Fig 8D–I.

Inspection of the dataset revealed that apical Müller glia form a dense population of thin tubular processes between photoreceptor inner segments, broadly distributed across the retinal plane and progressively decreasing in density with distance from the OLM (Fig 8B,C; Figure S10A–C,E,G). Cone-centered reconstructions further showed that multiple glial processes associate with each cone from the OLM to the ciliary region, with some extending distally along the cone outer segment. In the cone-centered subvolume, sparse RPE processes approached the cone from the apical side and capped the outer segment through multiple contact points. The longest RPE process extended basally to contact both the cone inner segment and a Müller glial process adjacent to the connecting cilium (Fig 8D–I; Figure S10F,H–J). Thus, although less elaborate than in zebrafish, the mouse subretinal space also displays a cone-associated organization in which glial and RPE processes converge around cones.

## Discussion

The subretinal space is becoming increasingly relevant in a clinical context. Drusenoid deposits can accumulate here in age-related macular degeneration [50], and the site is targeted for drug delivery and cell transplantation in potential treatments for retinal diseases [4]. Physiologically, the subretinal space is a tightly packed interface between photoreceptors and the RPE, but can expand in cases of retinal detachment, eventually leading to photoreceptor death if not rectified. It is therefore of critical importance to understand the architecture of the subretinal space and communication between its components.

Here, we provide a comprehensive view of the subretinal space in two well-established vertebrate models, zebrafish and mouse, using FIB-SEM complemented by confocal microscopy and we describe several unexpected findings discussed below. Notably, previous work presenting a serial block-face scanning electron microscopy (SBF-SEM) reconstruction of the human subretinal space reported that an incredible 38% of its volume is occupied by interphotoreceptor matrix [51], a specialized extracellular compartment rich in components mediating signaling and adhesion [52]. Similar proportions were reported in human transmission electron microscopy (TEM) data, with photoreceptors sparsely populating the subretinal space [53]. Still, apart from occasional tears in both datasets (most evident around the tip of the cone outer segments, see Fig. 8B; Figure S7, Figure S8), we did not observe any analogous gaps between cellular structures in our data. Our results are more comparable to FIB-SEM obtained in the mouse retina [54], images of larval zebrafish made with SBF-SEM [55], as well as to standard TEM and bright-field images of the zebrafish retina [56–58]. The discrepancy might come from variations in fixation protocols, delays in fixation of human tissue, or biological variation between species. Nevertheless, the gap between human photoreceptor outer segments is not apparent in live adaptive optics imaging [59, 60].

### Glia–RPE interaction is conserved between mouse and zebrafish

In this study, we characterize an intriguing cone-associated interface between the long apical processes of Müller glia and the RPE. Although this relationship was first reported in the frog over sixty years ago [16], the discovery has since been neglected by the field. Building on our previous observation of Müller glia–RPE overlap surrounding zebrafish UVS cones [17], we here provide a detailed morphological analysis of this interaction and, crucially, confirm its presence in a mammalian model.

While the interactions between RPE and glial processes around cone outer segments are a conserved feature, there is some divergence in organization of the subretinal space between zebrafish and mouse. In zebrafish, we show each UVS cone is intimately surrounded by 4–5 pairs of aligned Müller glial and RPE processes, stretching across the entire UVS cone outer segment. Long apical glial processes sprout from the OLM as isolated, thin structures but expand in cross-sectional area 10 times towards the tip, adopting an organization resembling bundled crayons. A similar appearance for glial processes has been reported in other species; however, they seem to emerge from the OLM already bundled [61–63]. In the mouse retina, Müller glial apical processes are thin tubular structures of approximately 100 nm in diameter that remain relatively constant along their length. Near the base of photoreceptor inner segment, they cluster at junctions between three neighbouring photoreceptors, where several processes run together as dense bundles. Only a sparse subset of Müller glial processes extends distally enough to reach and closely contact cone outer segments. The RPE reaches deep into the zebrafish subretinal tissue, forming a cap-like structure around a UVS cone outer segment tip and continuing as thin processes terminating close to the OLM. In the mouse retina, the RPE also provided substantial coverage of cone outer segment, though most RPE processes did not reach the inner segment. The ultrastructure of the RPE component is much less organized, with membrane indicative of ER filling the processes, as was also previously reported in human and rat electron microscopy datasets [51, 64]. Despite being often referred to as “microvilli” [9], long RPE processes appear more villous in their morphology.

### Establishment and characteristics of the glia–RPE interface

Our findings establish a comprehensive developmental timeline for the emergence of overlap between Müller glial and RPE processes around cone photoreceptors in the zebrafish and mouse retinas. In both species, the interaction occurs after outer segment formation and around the onset of visual function. While the two cell types extend simultaneously into the expanding subretinal space in zebrafish, mouse Müller glia are the first to arrange around developing cones, whereas RPE processes form later. Considering this timeline, glia–RPE contacts seem to be dispensable for the initial outer segment growth, but likely become required as the photoreceptor layer matures, increasing dependence on the supporting cells.

The gradual outgrowth of the processes and their morphology raise the question about the mechanistic framework at play. Interestingly, we observed the localization of pERM to apical Müller glia in both mouse and zebrafish. ERM proteins act as cross-linkers between the plasma membrane and actin filaments, playing a key role in microvillar formation and cell adhesion [65, 66]. In zebrafish, the localization of pERM in fluorescent micrographs is restricted to the most apical portion and corresponds to the position of the thickest part of the glial process in our FIB-SEM data, indicating pERM as a potential regulator of the bundled morphology. This is supported by the notion that both pERM expression and the bundled morphology appear more proximally in the mouse retina. In addition, ERM proteins were shown to interact *in vitro* with CD44 [66, 67], a Müller glial marker and a transmembrane hyaluronan receptor, abundant in glial apical processes in mouse [68]. As hyaluronic acid (HA) is a major component of the interphotoreceptor matrix [52], the ERM-CD44-HA axis is likely anchoring Müller glia within the subretinal space. Though CD44 is expressed in the zebrafish retina [69], its role there and its presence in Müller glia remain to be elucidated.

As previously shown, Crb2(a) is likely one of the components mediating the adhesion between apical Müller glial processes and photoreceptor inner segments [32]. Still, further research is necessary to determine the exact mechanism of the glia–RPE coupling. Another protein, junctional adhesion molecule C (JAM-C), was previously reported to localize to both Müller glial and RPE apical processes in the mouse retina and to undergo homophilic interactions and would be an interesting candidate to explore in future studies [21, 70].

### Functional implications of the glia–RPE contact

Our study reveals a previously underappreciated tripartite association between Müller glia, RPE, and cones in both zebrafish and mouse. While direct glia–RPE communication in the healthy retina is scarcely emphasized in the literature [16], such interactions are well documented in pathological conditions [71–74], and both cell types are known to influence one another through secreted factors [75–79]. This can be illustrated by the ability of pigment epithelium-derived factor (PEDF) to rescue degenerating Müller glia in cultured retinas after RPE removal [76]. Initially, we hypothesized that, in zebrafish, the cage-like enclosure formed by Müller glial and RPE processes around UVS cones was mainly required to locally support the demanding UVS cone metabolism. However, because this alignment persisted even after UVS cone loss, it is unlikely to function solely as a cone subtype-specific support system. Instead, glia–RPE interaction may reflect a more fundamental organizational principle of the subretinal space. Though tackling the exact role goes beyond the scope of this work, we consider a few possibilities below.

One possible role of this architecture is mechanical. Photoreceptor outer segments occupy much of the subretinal space while connected to the rest of the cell body via a narrow connecting cilium, creating a structurally vulnerable region. Müller glial cells were previously shown to physically reinforce the retinal layers in zebrafish [35], and our data suggest that apical RPE processes may complement this support, particularly in regions not fully spanned by Müller glia.

Another likely role for this structure is metabolic. The outer retina is a specialized metabolic unit, in which photoreceptor homeostasis depends on activities of both Müller glia and RPE [10, 11]. In this context, it is reasonable to suggest that the extension of both cell types into the subretinal space may facilitate the local transport of ions and nutrients as well as reuptake of metabolic waste products. The outer retina is avascular and relies on the choroid and the RPE for glucose delivery, while photoreceptors are the primary glucose consumers [80]. The expression of the glucose transporter, GLUT1, in both Müller glial and RPE apical processes [81], together with their close apposition within the subretinal space, raises the possibility that these compartments cooperate in local glucose transport within the subretinal space. Similarly, the presence of abundant ER in RPE apical processes, together with whorled multilamellar vesicles at Müller glial apical endfeet, is consistent with a metabolically active interface specialized for membrane handling, exchange, and possibly degradation. Such an extended contact zone could also support local delivery of RPE-derived trophic signals [76], and help buffer oxidative stress generated by the high metabolic activity of cones as well as their chronic exposure to light.

A third possibility is that this specialized interface contributes to retinoid recycling [82, 83]. Cone chromophore regeneration is supported by both Müller glia and RPE, and key components of the visual cycle are present in apical processes in both mouse and zebrafish. Rpe65, a key enzyme of the visual cycle, is present in the apical RPE in zebrafish and shows an expression pattern similar to that detected by the zpr-2 antibody [84]. In mouse, the visual cycle enzymes, including RPE65, LRAT, and RDH5 are restricted to the RPE cell body and do not extend into the apical processes [20]. However, the retinoid carrier protein CRALBP localizes to Müller glial and RPE apical processes of both mouse and zebrafish, potentially via an EBP50-Ezrin-dependent anchoring mechanism [19, 85]. At least in the RPE, most of the visual cycle enzymes are ER-associated [86]. Müller glia also contain ER in their apical domain, where local Ca^2+^ sensors trigger calcium influx following depletion of ER stores [87]. In this context, it is worth noting that the bundled crayon-like arrangement of glial processes we observed in zebrafish using FIB-SEM is reminiscent of smooth ER organized in hexagonal symmetry, known as a “crystalloid” ER [88, 89]. While this comparison is purely morphological, it is compatible with the possibility that the apical Müller glial and RPE compartments contain specialized ER that could support local retinoid recycling or other membrane associated functions. Although this interpretation remains speculative, the extensive overlap between glial and RPE apical territories raises the possibility that this interface acts as a functional junction between the two cone-supporting visual cycles.

In summary, our findings support a revised model of the subretinal architecture in which photoreceptors, particularly cones, reside within a structured and cooperative environment rather than a simple extracellular gap. Understanding the role of this newly recognized glial-epithelial interaction may reveal new principles of photoreceptor support and open new therapeutic avenues for retinal disease.

## Acknowledgments

We acknowledge the employees of North Campus Animal Services, University of Alberta and employees of the IRCM mouse facility for providing animal care. We are also very grateful to Drs.: Flavio Zolessi for sending us fixed *Tg(gfap:GFP)* fish, Andrew Simmonds for providing mounting medium and access to a Dragonfly-compatible computer, Sarah Hughes for access to the LSM700 microscope, Qiumin Tan for access to the Leica CM1520 cryostat, Hilmar Strickfaden for his support with FIB-SEM, Tom Hobman for sharing the 9E10 anti-Myc antibody, Ted Allison and Paul Chrystal for sharing the Tol2kit plasmids and crystal zebrafish line, Ryan McDonald for sharing *gfap*:GFP construct, Edan Foley for the Tol2 transposase mRNA, Craig Mandato and Hojatollah Vali at the Facility for Electron microscopy research at McGill University and Dominic Filion and Mathew Duguay at the IRCM microscopy core.

---

**Figure S1.** (A) A schematic drawing showing the retinal layers. RPE—retinal pigment epithelium, PRC—photoreceptor, OLM—outer limiting membrane, ONL—outer nuclear layer, OPL—outer plexiform layer, OS—outer segments, INL—inner nuclear layer, IPL—inner plexiform layer, GCL—ganglion cell layer. (B) A schematic depiction of the zebrafish photoreceptor layer and its organization, highlighting various proteins and their known position. (C–C”) Colocalization between actin labeled with phalloidin (cyan) and long apical glial processes (*gfap*:GFP; GFP—green) above the OLM. (D–D”) Colocalization of Crb2a labeled with zs-4 antibody (yellow) and long apical glial processes (*gfap*:GFP; GFP—green) above the OLM.

**Figure S2.** (A) *Tg(gfap:GFP)* (GFP—green) 1 mpf retina labeled with zrf-1 (magenta), n=3. (B) 1 mpf retina labeled with anti-GS antibody (green) and pERM (magenta), n=3. (C) An example of the injection efficiency and expression of *gfap*:GFP in the eye. Arrow points at the Müller glial processes above the OLM. Note that we noticed weak expression in isolated photoreceptors (asterisk). (D,E) Images of *Tg(hsp:act-myc)* 3.5 mpf zebrafish retina four hours after heat shock labeled with phalloidin (cyan) and anti-Myc (red) antibody, number of fish n=6. (D) Dark-adapted retina; arrowhead highlights a thick actin bundle around a UVS cone outer segment (OS). (E) Light-adapted retina. (F,G) Images of control *Tg(hsp:act-myc)* 3.5 mpf zebrafish retina labeled with phalloidin (cyan) and anti-Myc (red) antibody, dark-adapted (F) and light-adapted (G). Number of fish n=3. (H) Retinal micrograph of a 1 mpf Tg(sws1:GFP) zebrafish (GFP—yellow) stained with phalloidin (cyan) and pERM (red); only a few UVS cones remain due to a *tbx2a/b* CRISPR mosaic mutagenesis. (I) Images of a 1 mpf *Tg(sws1:GFP) tbx2a/b* UV+ crispant zebrafish (GFP—yellow) stained with zpr-2 (magenta). Arrowheads point at the RPE cap around the UV OS tip. (J–L) Images of a 1 mpf *Tg(sws1:GFP)* zebrafish (GFP—yellow) stained with phalloidin (cyan) and UV opsin antibody (yellow) to complement the GFP. Note the occasional UV opsin signal in the photoreceptor layer of crispants lacking UVS cones (L arrowheads). Scale bars: 20 μm (A,B,D–H, J–L), 50 μm (C), 10 μm (I).

**Figure S3. Measuring RPE presence at the OLM.** (A–C, E’) The graphs in which an area under the curve was measured, with the minimal threshold indicated (5000). (D) An example showing how the area around the OLM was measured. (E) Graph showing the intensity of the phalloidin signal measured at the OLM. Statistics: Welch’s ANOVA test, Dunnett’s T3 multiple comp. test, median is shown; n=6 (control), n=8 (*tbx2a/b* UV+), n=9 (*tbx2a/b* UV-).

**Figure S4. Endogenous Müller glial process, actin, and cone morphology and protein localization.** (A) Immunolabeled images of wild-type mouse retinal cross-sections at the indicated stages of post-natal development. Müller glial apical processes (CD44, green) and actin-rich apical processes (pERM, magenta) are shown. Dashed line indicates OLM. Scale bar: 3 μm. (B) Same as (A) but lower magnification to show full cone morphology (ARR3, magenta) and F-actin (phalloidin, green). Scale bar: 10 μm.

**Figure S5. Electroporated RPE cells exhibit normal morphology and extend long apical processes.** (A) En face epi-fluorescence images of a whole-mounted adult mouse RPE (left) or corresponding neural retina (right) electroporated with mNeonGreen-CAAX construct. Images illustrate robust RPE electroporation and the extension of apical RPE processes into the neural retinal layer. Scale bar: 300 μm. (B) Epifluorescence images of adult mouse retinal cross sections showing three examples of a single RPE cell electroporated with mNeonGreen-CAAX. Dashed line indicates OLM. Scale bar: 10 μm. (C) En face, high magnification image of whole-mounted adult mouse RPE showing electroporated RPE cells retain normal epithelial cell morphology. Scale bar: 40 μm. (D) 3D projection of three electroporated RPE cells (mNeonGreen, cyan) superimposed with phalloidin (magenta, right) to illustrate normal tight junction morphology. (E) En face epifluorescence and immunofluorescence images (top) and corresponding 3D reconstruction (bottom) of electroporated RPE cell (cyan), phalloidin (magenta), and p-ERM showing normal RPE cell morphology and lateral extensions (arrows). Scale bar: 10 μm. (F) Projection of en face acquisition in XZ plane showing mNeonGreen decoration of both apical and basal RPE membrane. Scale bar: 3 μm. (G) 3D reconstruction and zoom in of the longest lateral process shown in white dashed box in (E). (H) Epifluorescence and immunofluorescence images of an adult mouse retinal cross section showing a long apical extension from an electroporated RPE cell that makes contact with the OLM (black dashed line). Scale bar: 2 μm.

**Figure S6. No obvious differences in electroporated RPE cells in dark-adapted mice compared to light-adapted mice.** (A) Epifluorescence and immunofluorescence images of an adult mouse retinal cross section following dark adaptation. Dashed line indicates OLM. Scale bar: 5 μm. Zoomed in images show presence of pERM in the proximal processes, and the absence of pERM more distally. Scale bar: 1 μm. (B) Same as (A) showing a light-adapted adult mouse retinal cross section.

**Figure S7. Block surface and region targeted for zebrafish FIB-SEM imaging.**

**Figure S8 Morphology of apical Müller glia and RPE in zebrafish at different levels.** (A-G) Annotated FIB-SEM slices with segmented ROIs visible: UVS inner segment—blue, UVS outer segment—yellow, Müller glia—green, RPE—magenta. Images are organized sequentially from UVS inner segment at the OLM to the level above the UVS outer segment tip. (C) Note how the base of the UVS outer segment is surrounded by an inner segment sheet, which has not divided into microvilli yet. Arrowhead points at the accessory outer segment. (G) Arrowheads show pigment granules within the RPE extension. (H) Maximum intensity projection of a confocal stack of 1 mpf zebrafish photoreceptor layer stained with anti-calreticulin antibody, phalloidin (cyan), and zpr-2 (magenta). Arrowheads highlight localization of calreticulin to the apical RPE processes.

**Figure S9. Block surface and region targeted for mouse FIB-SEM imaging.**

**Figure S10. Müller glial and RPE apical processes form a tubular network between photoreceptors and contact cones in the mouse subretinal space.** (A) Subvolume from a FIB-SEM acquisition of the mouse subretinal region. Orthogonal projections are shown together with the original en face images acquired at different depths. Müller glial (MG) processes are identified as individual round structures (green dots). Scale bar: 4 μm. (B) FIB-SEM images acquired at increasing distances from the OLM within the subretinal space. The density of MG processes (green) decreases with distance from the OLM. (C) Quantification of MG process density as a function of distance from the OLM. Dashed lines represent the average depth of the photoreceptor inner segment-to-outer segment transition zone (TZ). (D) FIB-SEM micrograph showing optically dense MG-photoreceptor junctions forming the OLM. White arrowheads point to whorl-like structures inside vesicles at the apical MG end feet. Scale bar: 1 um. (E) FIB-SEM micrograph showing MG cell processes located between photoreceptor inner segments (IS, green arrows). Note that MG processes are typically located at the junction of three photoreceptor ISs. Scale bar: 1 μm. (F) High magnification FIB-SEM micrograph at the level of the cone outer segment (OS) showing presumptive RPE apical processes (magenta arrowheads). Artifactual delamination of cone OS was observed presumably due to fixation (*). Black arrows indicate vesicular structures associated with the delaminated cone membranes. (G) 3D reconstruction of cone subcompartments and MG apical processes from the entire FIB-SEM dataset. Whole volume AI-mediated (left) and manual reconstruction of cone-associated processes (right). The AI-mediated reconstruction shows a homogenous distribution of MG cell processes across the retinal plane, whereas manual reconstruction of individual processes in the vicinity of cones highlights their interactions with cones. Green, MG processes; yellow, cone OS; cyan, cone IS; red, connecting cilium; blue, axoneme. (H) High-magnification manual 3D reconstruction of a cone at the ciliary region (left) and transverse XY projection (middle), showing a MG process (arrows) adjacent to the cone OS axoneme (blue). Right panel: high-magnification en face images acquired at increasing distal distances (z) from the base of the axoneme illustrate the spatial relationship between the MG process (arrows) and the cone OS (yellow), identified by its electron-dense membrane. (I) Quantification of MG process diameter at different positions along cone length (n=8 cones). Each dot represents one MG process. (J) Quantification of the number of MG cell processes associated with each cone at different positions along cone length. All cones were associated with processes up to the ciliary region (black dots), whereas some cones lacked associated processes along their outer segments (grey dots).

**Video S1.** Slice and view of 3D signal inferred from en face imaging stacks of CD44 (green) and ARR3 (magenta) immunostaining from a whole-mounted mouse retina. 3D surface reconstructions of cone outer segment (yellow), cone inner segment (cyan), Müller glial processes (green) are shown.

**Video S2.** Slice and view of 3D signal inferred from en face imaging stacks of CD44 (green) and S-opsin (magenta) immunostaining following mScarlet3-CAAX (cyan) RPE electroporation and retinal whole mount. 3D surface reconstructions of cone outer segment (magenta), electroporated RPE process (cyan), and Müller glial process (green) are shown.

**Video S3.** 3D reconstruction of a UVS photoreceptor and surrounding Müller glia and RPE in a zebrafish retina.

**Video S4.** Scroll through all the FIB-SEM sections with segmented ROIs visible. Four classes were painted around a single UVS cone: UVS inner segment—blue, UVS outer segment—yellow, Müller glia—green, RPE—magenta.

**Video S5.** Slice and view through the whole mouse FIB-SEM outer retina volume. 3D reconstruction of cone outer segment (yellow), inner segment (cyan), connecting cilium (red), axoneme (blue). Müller glial apical processes associated with each cone were manually reconstructed (green lines), while the global glial network was reconstructed using AI (light green).

**Video S6.** Slice and view through the cropped mouse FIB-SEM outer retina volume. 3D reconstruction of cone outer segment (yellow), inner segment (cyan), connecting cilium (red), axoneme (blue), putative RPE processes (magenta), Müller glial processes (green). Final frames of video shows RPE and glial structures generated through AI pixel classification tool.

## References

1. Richard W. Young. The renewal of photoreceptor cell outer segments. The Journal of Cell Biology, 33(1):61–72, April 1967. ISSN 0021-9525. doi: 10.1083/jcb.33.1.61.

2. Matthew M. LaVail. Rod outer segment disk shedding in rat retina: relationship to cyclic lighting. Science, 194(4269):1071–1074, December 1976. ISSN 1095-9203. doi: 10.1126/science.982063.

3. Marion Sangster Eckmiller. Morphogenesis and renewal of cone outer segments. Progress in Retinal and Eye Research, 16(3):401–441, July 1997. ISSN 1350-9462. doi: 10.1016/s1350-9462(96)00026-2.

4. Nina Buus Sørensen. Subretinal surgery: functional and histological consequences of entry into the subretinal space. Acta Ophthalmologica, 97(A114):1–23, November 2019. ISSN 1755-3768. doi: 10.1111/aos.14249.

5. Radgonde Amer, Hilal Nalcı, and Nilüfer Yalçındağ. Exudative retinal detachment. Survey of Ophthalmology, 62(6):723–769, November 2017. ISSN 0039-6257. doi: 10.1016/j.survophthal.2017.05.001.

6. S. Omri, B. Omri, M. Savoldelli, L. Jonet, B. Thillaye-Goldenberg, G. Thuret, P. Gain, J. C. Jeanny, P. Crisanti, and F. Behar-Cohen. The outer limiting membrane (OLM) revisited: clinical implications. Clinical Ophthalmology, page 183, March 2010. ISSN 1177-5483. doi: 10.2147/opth.s5901.

7. Jonathan B. Lin, Raja Narayanan, Elise Philippakis, Yoshihiro Yonekawa, and Rajendra S. Apte. Retinal detachment. Nature Reviews Disease Primers, 10(1), March 2024. ISSN 2056-676X. doi: 10.1038/s41572-024-00501-5.

8. Isabela Martins Melo, Tianwei Ellen Zhou, Flavia Nagel, Nikhil S. Patil, Fathima Afira Faleel, Marko Popovic, and Rajeev H. Muni. Histological changes in retinal detachment: A systematic review for the clinician. Survey of Ophthalmology, 69(1):85–92, January 2024. ISSN 0039-6257. doi: 10.1016/j.survophthal.2023.08.001.

9. Olaf Strauss. The retinal pigment epithelium in visual function. Physiological Reviews, 85(3):845–881, July 2005. ISSN 1522-1210. doi: 10.1152/physrev.00021.2004.

10. Mark A. Kanow, Michelle M. Giarmarco, Connor S. R. Jankowski, Kristine Tsantilas, Abbi L. Engel, Jianhai Du, Jonathan D. Linton, Christopher C. Farnsworth, Stephanie R. Sloat, Austin Rountree, Ian R. Sweet, Ken J. Lindsay, Edward D. Parker, Susan E. Brockerhoff, Martin Sadilek, Jennifer R. Chao, and James B. Hurley. Biochemical adaptations of the retina and retinal pigment epithelium support a metabolic ecosystem in the vertebrate eye. eLife, 6, September 2017. ISSN 2050-084X. doi: 10.7554/elife.28899.

11. Joseph Hanna, Luke Ajay David, Yacine Touahri, Taylor Fleming, Robert A. Screaton, and Carol Schuurmans. Beyond genetics: the role of metabolism in photoreceptor survival, development and repair. Frontiers in Cell and Developmental Biology, 10, May 2022. ISSN 2296-634X. doi: 10.3389/fcell.2022.887764.

12. R.h. Steinberg, I. Wood, and M.j. Hogan. Pigment epithelial ensheathment and phagocytosis of extrafoveal cones in human retina. Philosophical Transactions of the Royal Society of London. B, Biological Sciences, 277(958):459–471, March 1977. ISSN 2054-0280. doi: 10.1098/rstb.1977.0028.

13. R. H. Steinberg and I. Wood. Pigment epithelial cell ensheathment of cone outer segments in the retina of the domestic cat. Proceedings of the Royal Society of London. Series B. Biological Sciences, 187(1089):461–478, November 1974. ISSN 2053-9193. doi: 10.1098/rspb.1974.0088.

14. Steven K. Fisher and Roy H. Steinberg. Origin and organization of pigment epithelial apical projections to cones in cat retina. Journal of Comparative Neurology, 206(2):131–145, April 1982. ISSN 1096-9861. doi: 10.1002/cne.902060204.

15. Bruce A. Pfeffer and Steven K. Fisher. Development of retinal pigment epithelial surface structures ensheathing cone outer segments in the cat. Journal of Ultrastructure Research, 76(2):158–172, August 1981. ISSN 0022-5320. doi: 10.1016/s0022-5320(81)80014-7.

16. Sven Erik G. Nilsson. An electron microscopic classification of the retinal receptors of the leopard frog (Rana pipiens). Journal of Ultrastructure Research, 10(5–6):390–416, June 1964. ISSN 0022-5320. doi: 10.1016/s0022-5320(64)80018-6.

17. Maria Sharkova, Gonzalo Aparicio, Constantin Mouzaaber, Flavio R. Zolessi, and Jennifer C. Hocking. Photoreceptor calyceal processes accompany the developing outer segment, adopting a stable length despite a dynamic core. Journal of Cell Science, 137(7), April 2024. ISSN 1477-9137. doi: 10.1242/jcs.261721.

18. Andreas Reichenbach and Andreas Bringmann. Müller cells in the healthy and diseased retina. Springer New York, 2010. ISBN 9781441916723. doi: 10.1007/978-1-4419-1672-3.

19. Maria Nawrot, Karen West, Jing Huang, Daniel E. Possin, Anthony Bretscher, John W. Crabb, and John C. Saari. Cellular retinaldehyde-binding protein interacts with ERM-binding phosphoprotein 50 in retinal pigment epithelium. Investigative Ophthalmology & Visual Science, 45(2):393, February 2004. ISSN 1552-5783. doi: 10.1167/iovs.03-0989.

20. Jing Huang, Daniel E. Possin, and John C. Saari. Localizations of visual cycle components in retinal pigment epithelium. Molecular Vision, 15:223–234, 2009. URL http://www.molvis.org/molvis/v15/a22.

21. Lauren L. Daniele, Ralf H. Adams, Diane E. Durante, Edward N. Pugh, and Nancy J. Philp. Novel distribution of junctional adhesion molecule-C in the neural retina and retinal pigment epithelium. Journal of Comparative Neurology, 505(2):166–176, September 2007. ISSN 1096-9861. doi: 10.1002/cne.21489.

22. Masaki Takechi, Takanori Hamaoka, and Shoji Kawamura. Fluorescence visualization of ultraviolet-sensitive cone photoreceptor development in living zebrafish. FEBS Letters, 553(1–2):90–94, September 2003. ISSN 1873-3468. doi: 10.1016/s0014-5793(03)00977-3.

23. Rebecca L. Bernardos and Pamela A. Raymond. GFAP transgenic zebrafish. Gene Expression Patterns, 6(8):1007–1013, October 2006. ISSN 1567-133X. doi: 10.1016/j.modgep.2006.04.006.

24. Spencer Balay. Cryptochrome expression in the zebrafish retina: potential implications for magnetoreception. PhD thesis, University of Alberta, Edmonton, Canada, 2018.

25. Paride Antinucci and Robert Hindges. A crystal-clear zebrafish for in vivo imaging. Scientific Reports, 6(1), July 2016. ISSN 2045-2322. doi: 10.1038/srep29490.

26. Juan M. Angueyra, Vincent P. Kunze, Laura K. Patak, Hailey Kim, Katie Kindt, and Wei Li. Transcription factors underlying photoreceptor diversity. eLife, 12, February 2023. ISSN 2050-084X. doi: 10.7554/elife.81579.

27. Kazuyuki Hoshijima, Michael J. Jurynec, Dana Klatt Shaw, Ashley M. Jacobi, Mark A. Behlke, and David Jonah Grunwald. Highly efficient CRISPR-Cas9-based methods for generating deletion mutations and F0 embryos that lack gene function in zebrafish. Developmental Cell, 51(5):645–657.e4, December 2019. ISSN 1534-5807. doi: 10.1016/j.devcel.2019.10.004.

28. Ye Liu, Lily Ng, Hong Liu, Heike Heuer, and Douglas Forrest. Cone photoreceptor differentiation regulated by thyroid hormone transporter MCT8 in the retinal pigment epithelium. Proceedings of the National Academy of Sciences, 121(30), July 2024. ISSN 1091-6490. doi: 10.1073/pnas.2402560121.

29. Christiana J. Johnson, Lennart Berglin, Micah A. Chrenek, T. M. Redmond, Jeffrey H. Boatright, and John M. Nickerson. Technical brief: subretinal injection and electroporation into adult mouse eyes. Molecular vision, 14:2211–2226, 2008. ISSN 1090-0535.

30. Johannes Schindelin, Ignacio Arganda-Carreras, Erwin Frise, Verena Kaynig, Mark Longair, Tobias Pietzsch, Stephan Preibisch, Curtis Rueden, Stephan Saalfeld, Benjamin Schmid, Jean-Yves Tinevez, Daniel James White, Volker Hartenstein, Kevin Eliceiri, Pavel Tomancak, and Albert Cardona. Fiji: an open-source platform for biological-image analysis. Nature Methods, 9(7):676–682, June 2012. ISSN 1548-7105. doi: 10.1038/nmeth.2019.

31. Tiago Ferreira, Kota Miura, Bitdeli Chef, and Jan Eglinger. Scripts: BAR 1.1.6, 2015.

32. Jian Zou, Xiaolei Wang, and Xiangyun Wei. Crb apical polarity proteins maintain zebrafish retinal cone mosaics via intercellular binding of their extracellular domains. Developmental Cell, 22(6): 1261–1274, June 2012. ISSN 1534-5807. doi: 10.1016/j.devcel.2012.03.007.

33. Chuanyu Guo, Ciana Deveau, Cen Zhang, Ralph Nelson, and Xiangyun Wei. Zebrafish Crb1, localizing uniquely to the cell membranes around cone photoreceptor axonemes, alleviates light damage to photoreceptors and modulates cones’ light responsiveness. The Journal of Neuroscience, 40(37): 7065–7079, August 2020. ISSN 1529-2401. doi: 10.1523/jneurosci.0497-20.2020.

34. Maria Toms, Thomas Burgoyne, Dhani Tracey-White, Rose Richardson, Adam M. Dubis, Andrew R. Webster, Clare Futter, and Mariya Moosajee. Phagosomal and mitochondrial alterations in RPE may contribute to KCNJ13 retinopathy. Scientific Reports, 9(1), March 2019. ISSN 2045-2322. doi: 10.1038/s41598-019-40507-8.

35. Ryan B. MacDonald, Owen Randlett, Julia Oswald, Takeshi Yoshimatsu, Kristian Franze, and William A. Harris. Müller glia provide essential tensile strength to the developing retina. Journal of Cell Biology, 210(7):1075–1083, September 2015. ISSN 1540-8140. doi: 10.1083/jcb.201503115.

36. Stefan Kustermann, Herbert Hildebrandt, Sylvia Bolz, Katja Dengler, and Konrad Kohler. Genesis of rods in the zebrafish retina occurs in a microenvironment provided by polysialic acid-expressing Müller glia. Journal of Comparative Neurology, 518(5):636–646, December 2009. ISSN 1096-9861. doi: 10.1002/cne.22232.

37. A. Nicol San Juan, Allison D. N. N. Tran, Debarati Hore, Linnea M. Kriese, Natalie H. N. Van Engelen, Maria Sharkova, Nicole C. L. Noel, Carmanah Hunter, Olga Spencer Carreira, Jennifer C. Hocking, Lisa M. Willis, and Brittany J. Carr. Polysialic acid is a versatile marker for retinal Müller glia in common vertebrate model organisms and systems. September 2025. doi: 10.1101/2025.09.04.672712.

38. Dirk Höfer and Detlev Drenckhahn. Molecular heterogeneity of the actin filament cytoskeleton associated with microvilli of photoreceptors, Müller’s glial cells and pigment epithelial cells of the retina. Histochemistry, 99(1):29–35, January 1993. ISSN 0301-5564. doi: 10.1007/bf00268017.

39. Tero Kivelä, Juha Jääskeläine, Antti Vaheri, and Olli Carpén. Ezrin, a membrane-organizing protein, as a polarization marker of the retinal pigment epithelium in vertebrates. Cell and Tissue Research, 301(2):217–223, July 2000. ISSN 1432-0878. doi: 10.1007/s004410000225.

40. Pavan Vedula, Marie E. Fina, Brent A. Bell, Sergei S. Nikonov, Anna Kashina, and Dawei W. Dong. -actin is essential for structural integrity and physiological function of the retina. March 2023. doi: 10.1101/2023.03.27.534392.

41. Nicholas J. Hanovice, Lyndsay L. Leach, Kayleigh Slater, Ana E. Gabriel, Dwight Romanovicz, Enhua Shao, Ross Collery, Edward A. Burton, Kira L. Lathrop, Brian A. Link, and Jeffrey M. Gross. Regeneration of the zebrafish retinal pigment epithelium after widespread genetic ablation. PLOS Genetics, 15(1):e1007939, January 2019. ISSN 1553-7404. doi: 10.1371/journal.pgen.1007939.

42. Stephen Yazulla and Keith M. Studholme. Neurochemical anatomy of the zebrafish retina as determined by immunocytochemistry. Journal of Neurocytology, 30(7):551–592, 2001. ISSN 1573-7381. doi: 10.1023/a:1016512617484.

43. Beth Burnside and Barbara Nagle. Chapter 3. retinomotor movements of photoreceptors and retinal pigment epithelium: Mechanisms and regulation. Progress in Retinal Research, 2:67–109, January 1983. ISSN 0278-4327. doi: 10.1016/0278-4327(83)90004-4.

44. Michael H. Chaitin, Helen S. Wortham, and Anne-Marie Brun-Zinkernagel. Immunocytochemical localization of CD44 in the mouse retina. Experimental Eye Research, 58(3):359–365, March 1994. ISSN 0014-4835. doi: 10.1006/exer.1994.1026.

45. Pengfei Zhang, Bradley Shibata, Gabriel Peinado, Robert J. Zawadzki, Paul FitzGerald, and Edward N. Pugh. Measurement of diurnal variation in rod outer segment length in vivo in mice with the oct optoretinogram. Investigative Opthalmology amp; Visual Science, 61(3):9, March 2020. ISSN 1552-5783. doi: 10.1167/iovs.61.3.9.

46. Zhenqing Zhou, Frans Vinberg, Frank Schottler, Teresa A Doggett, Vladimir J Kefalov, and Thomas A Ferguson. Autophagy supports color vision. Autophagy, 11(10):1821–1832, October 2015. ISSN 1554-8635. doi: 10.1080/15548627.2015.1084456.

47. Ellen A. Schmitt and John E. Dowling. Early retinal development in the zebrafish, Danio rerio: Light and electron microscopic analyses. The Journal of Comparative Neurology, 404(4):515–536, February 1999. ISSN 1096-9861. doi: 10.1002/(sici)1096-9861(19990222)404:4<515::aid-cne8>3.0.co;2-a.

48. Cátia Crespo and Elisabeth Knust. Characterisation of maturation of photoreceptor cell sub-types during zebrafish retinal development. Biology Open, January 2018. ISSN 2046-6390. doi: 10.1242/bio.036632.

49. Michael Housset, Dominic Filion, Nelson Cortes, Hojatollah Vali, Craig A. Mandato, Christian Casanova, and Michel Cayouette. Identification of a non-canonical planar cell polarity pathway triggered by light in the developing mouse retina. Developmental Cell, 60(3):447–458.e5, February 2025. ISSN 1534-5807. doi: 10.1016/j.devcel.2024.10.012.

50. Manuel Monge, Adriana Araya, and Lihteh Wu. Subretinal drusenoid deposits: An update. Taiwan Journal of Ophthalmology, 12(2):138–146, April 2022. ISSN 2211-5056. doi: 10.4103/tjo.tjo_18_22.

51. Maximilian Lindell, Deepayan Kar, Aleksandra Sedova, Yeon Jin Kim, Orin S. Packer, Ursula Schmidt-Erfurth, Kenneth R. Sloan, Mike Marsh, Dennis M. Dacey, Christine A. Curcio, and Andreas Pollreisz. Volumetric reconstruction of a human retinal pigment epithelial cell reveals specialized membranes and polarized distribution of organelles. Investigative Ophthalmology & Visual Science, 64(15):35, December 2023. ISSN 1552-5783. doi: 10.1167/iovs.64.15.35.

52. Makoto Ishikawa, Yu Sawada, and Takeshi Yoshitomi. Structure and function of the interphotoreceptor matrix surrounding retinal photoreceptor cells. Experimental Eye Research, 133:3–18, April 2015. ISSN 0014-4835. doi: 10.1016/j.exer.2015.02.017.

53. Tylor R. Lewis, Natalia V. Klementieva, Sebastien Phan, Carson M. Castillo, Keun-Young Kim, Lauren Y. Cao, Mark H. Ellisman, Vadim Y. Arshavsky, and Oleg Alekseev. Unique ultrastructural organization of human rod photoreceptors. Communications Biology, 8(1), January 2025. ISSN 2399-3642. doi: 10.1038/s42003-025-07473-6.

54. Sanjay Ch, Rayne R. Lim, Shermaine W. Y. Low, Deana G. Grant, Sam Patterson, Aparna Ramasubramanian, Ashish K. Gadicherla, and Shyam S. Chaurasia. A comprehensive overview of focused ion beam-scanning electron microscopy (FIB-SEM) applications for the evaluation of outer retina. Frontiers in Cell and Developmental Biology, 13, May 2025. ISSN 2296-634X. doi: 10.3389/fcell.2025.1586029.

55. Rachel A. Hutto, Kaitlyn M. Rutter, Michelle M. Giarmarco, Edward D. Parker, Zachary S. Chambers, and Susan E. Brockerhoff. Cone photoreceptors transfer damaged mitochondria to Müller glia. Cell Reports, 42(2):112115, February 2023. ISSN 2211-1247. doi: 10.1016/j.celrep.2023.112115.

56. Ruxandra Bachmann-Gagescu, Ian G. Phelps, George Stearns, Brian A. Link, Susan E. Brockerhoff, Cecilia B. Moens, and Dan Doherty. The ciliopathy gene cc2d2a controls zebrafish photoreceptor outer segment development through a role in Rab8-dependent vesicle trafficking. Human Molecular Genetics, 20(20):4041–4055, August 2011. ISSN 0964-6906. doi: 10.1093/hmg/ddr332.

57. Nicole C. L. Noel, W. Ted Allison, Ian M. MacDonald, and Jennifer C. Hocking. Zebrafish and inherited photoreceptor disease: Models and insights. Progress in Retinal and Eye Research, 91: 101096, November 2022. ISSN 1350-9462. doi: 10.1016/j.preteyeres.2022.101096.

58. Fei Liu, Jiaxiang Chen, Shanshan Yu, Rakesh Kotapati Raghupathy, Xiliang Liu, Yayun Qin, Chang Li, Mi Huang, Shengjie Liao, Jiuxiang Wang, Jian Zou, Xinhua Shu, Zhaohui Tang, and Mugen Liu. Knockout of RP2 decreases GRK1 and rod transducin subunits and leads to photoreceptor degeneration in zebrafish. Human Molecular Genetics, 24(16):4648–4659, June 2015. ISSN 1460-2083. doi: 10.1093/hmg/ddv197.

59. Austin Roorda and David R. Williams. The arrangement of the three cone classes in the living human eye. Nature, 397(6719):520–522, February 1999. ISSN 1476-4687. doi: 10.1038/17383.

60. Zhuolin Liu, Omer P. Kocaoglu, and Donald T. Miller. 3D imaging of retinal pigment epithelial cells in the living human retina. Investigative Ophthalmology & Visual Science, 57(9):OCT533, August 2016. ISSN 1552–5783. doi: 10.1167/iovs.16-19106.

61. Manfred Spitznas. Zur feinstruktur der sog. membrana limitans externa der menschlichen retina. Albrecht von Graefe’s Archive for Clinical and Experimental Ophthalmology, 180(1):44–56, May 1970. ISSN 1435-702X. doi: 10.1007/bf02384821.

62. Ingrid P. Meschede, Nicholas C. Ovenden, Miguel C. Seabra, Clare E. Futter, Marcela Votruba, Michael E. Cheetham, and Thomas Burgoyne. Symmetric arrangement of mitochondria: plasma membrane contacts between adjacent photoreceptor cells regulated by Opa1. Proceedings of the National Academy of Sciences, 117(27):15684–15693, June 2020. ISSN 1091–6490. doi: 10.1073/pnas.2000304117.

63. S. Uga and G. K. Smelser. Comparative study of the fine structure of retinal Müller cells in various vertebrates. Investigative Ophthalmology & Visual Science, 12:434–448, June 1973. ISSN 0020-9988.

64. John E. Dowling and I. R. Gibbons. The fine structure of the pigment epithelium in the albino rat. The Journal of Cell Biology, 14(3):459–474, September 1962. ISSN 0021-9525. doi: 10.1083/jcb.14.3.459.

65. Shigenobu Yonemura, Sachiko Tsukita, and Shoichiro Tsukita. Direct involvement of Ezrin/Radixin/Moesin (ERM)-binding membrane proteins in the organization of microvilli in collaboration with activated ERM proteins. The Journal of Cell Biology, 145(7):1497–1509, June 1999. ISSN 1540-8140. doi: 10.1083/jcb.145.7.1497.

66. Tomoyuki Mori, Ken Kitano, Shin-ichi Terawaki, Ryoko Maesaki, Yayoi Fukami, and Toshio Hakoshima. Structural basis for CD44 recognition by ERM proteins. Journal of Biological Chemistry, 283(43):29602–29612, October 2008. ISSN 0021-9258. doi: 10.1074/jbc.m803606200.

67. Shigenobu Yonemura, Motohiro Hirao, Yoshinori Doi, Nobuyuki Takahashi, Takahisa Kondo, Sachiko Tsukita, and Shoichiro Tsukita. Ezrin/Radixin/Moesin (ERM) proteins bind to a positively charged amino acid cluster in the juxta-membrane cytoplasmic domain of CD44, CD43, and ICAM-2. The Journal of Cell Biology, 140(4):885–895, February 1998. ISSN 1540-8140. doi: 10.1083/jcb.140.4.885.

68. Monika Ayten, Tobias Straub, Lew Kaplan, Stefanie M. Hauck, Antje Grosche, and Susanne F. Koch. CD44 signaling in Müller cells impacts photoreceptor function and survival in healthy and diseased retinas. Journal of Neuroinflammation, 21(1), August 2024. ISSN 1742-2094. doi: 10.1186/s12974-024-03175-8.

69. Sarah E. Grabinski, Dhwani Parsana, and Brian D. Perkins. Comparative analysis of transcriptional changes in zebrafish cep290 and bbs2 mutants by RNA-seq reveals upregulation of inflammatory and stress-related pathways. Frontiers in Molecular Neuroscience, 16, May 2023. ISSN 1662-5099. doi: 10.3389/fnmol.2023.1148840.

70. Sentot Santoso, Valeria V. Orlova, Kaimei Song, Ulrich J. Sachs, Cornelia L. Andrei-Selmer, and Triantafyllos Chavakis. The homophilic binding of junctional adhesion molecule-C mediates tumor cell-endothelial cell interactions. Journal of Biological Chemistry, 280(43):36326–36333, October 2005. ISSN 0021-9258. doi: 10.1074/jbc.m505059200.

71. Daniel Larbi, Alexander M. Rief, Seoyoung Kang, Shaoheng Chen, Khulan Batsuuri, Sabine Fuhrmann, Suresh Viswanathan, and Stefanie G. Wohl. Dicer loss in Müller glia leads to a defined sequence of pathological events beginning with cone dysfunction. Investigative Ophthalmology & Visual Science, 66(3):7, March 2025. ISSN 1552-5783. doi: 10.1167/iovs.66.3.7.

72. Dongfeng Cao, Belinda Leong, Jeffrey D. Messinger, Deepayan Kar, Thomas Ach, Lawrence A. Yannuzzi, K. Bailey Freund, and Christine A. Curcio. Hyperreflective foci, optical coherence tomography progression indicators in age-related macular degeneration, include transdifferentiated retinal pigment epithelium. Investigative Ophthalmology & Visual Science, 62(10):34, August 2021. ISSN 1552-5783. doi: 10.1167/iovs.62.10.34.

73. Rouel S. Roque and Ruth B. Caldwell. Müller cell changes precede vascularization of the pigment epithelium in the dystrophic rat retina. Glia, 3(6):464–475, January 1990. ISSN 1098-1136. doi: 10.1002/glia.440030605.

74. Mike Francke, Felix Makarov, Johannes Kacza, Johannes Seeger, Susanne Wendt, Ulrich Gärtner, Frank Faude, Peter Wiedemann, and Andreas Reichenbach. Retinal pigment epithelium melanin granules are phagocytozed by Müller glial cells in experimental retinal detachment. Journal of Neurocytology, 30(2):131–136, February 2001. ISSN 1573-7381. doi: 10.1023/a:1011987107034.

75. Mark K. Webster, Betty J. Barnett, Megan L. Stanchfield, Joshua R. Paris, Sarah E. Webster, Cynthia A. Cooley-Themm, Edward M. Levine, Deborah C. Otteson, and Cindy L. Linn. Stimulation of retinal pigment epithelium with an 7 nAChR agonist leads to Müller glia dependent neurogenesis in the adult mammalian retina. Investigative Ophthalmology & Visual Science, 60(2):570, February 2019. ISSN 1552-5783. doi: 10.1167/iovs.18-25722.

76. Monica M. Jablonski, Joyce Tombran-Tink, David A. Mrazek, and Alessandro Iannaccone. Pigment epithelium-derived factor supports normal Müller cell development and glutamine synthetase expression after removal of the retinal pigment epithelium. Glia, 35(1):14–25, May 2001. ISSN 1098-1136. doi: 10.1002/glia.1066.

77. C. David Jaynes, Harold J. Sheedlo, Neeraj Agarwal, Katherine O’ Rourke, and James E. Turner. Müller and retinal pigment epithelial (RPE) cell expression of NGFI-A and c-fos mRNA in response to medium conditioned by the RPE. Molecular Brain Research, 32(2):329–337, September 1995. ISSN 0169-328X. doi: 10.1016/0169-328x(95)00093-8.

78. Iwona Malgorzata Goczalik, Maik Raap, Michael Weick, Ivan Milenkovic, Jördis Heidmann, Volker Enzmann, Peter Wiedemann, Andreas Reichenbach, and Mike Francke. The activation of IL-8 receptors in cultured guinea pig Müller glial cells is modified by signals from retinal pigment epithelium. Journal of Neuroimmunology, 161(1–2):49–60, April 2005. ISSN 0165-5728. doi: 10.1016/j.jneuroim.2004.12.004.

79. Gisela Velez, Alexa R. Weingarden, Budd A. Tucker, Hetian Lei, Andrius Kazlauskas, and Michael J. Young. Retinal pigment epithelium and Müller progenitor cell interaction increase Müller progenitor cell expression of PDGFR and ability to induce proliferative vitreoretinopathy in a rabbit model. Stem Cells International, 2012:1–6, 2012. ISSN 1687-9678. doi: 10.1155/2012/106486.

80. Thierry Léveillard and José-Alain Sahel. Metabolic and redox signaling in the retina. Cellular and Molecular Life Sciences, 74(20):3649–3665, August 2016. ISSN 1420-9071. doi: 10.1007/s00018-016-2318-7.

81. L. Bergersen, E. Jóhannsson, M. L. Veruki, E. A. Nagelhus, A. Halestrap, O. M. Sejersted, and O. P. Ottersen. Cellular and subcellular expression of monocarboxylate transporters in the pigment epithelium and retina of the rat. Neuroscience, 90(1):319–331, February 1999. ISSN 0306-4522. doi: 10.1016/s0306-4522(98)00427-8.

82. Nathan L. Mata, Roxana A. Radu, Richard S. Clemmons, and Gabriel H. Travis. Isomerization and oxidation of vitamin A in cone-dominant retinas. Neuron, 36(1):69–80, September 2002. ISSN 0896-6273. doi: 10.1016/s0896-6273(02)00912-1.

83. Shinya Sato and Vladimir J. Kefalov. The retina-based visual cycle. Annual Review of Vision Science, 10(1):293–321, September 2024. ISSN 2374-4650. doi: 10.1146/annurev-vision-100820-083937.

84. Ryan MacDonald. Section of zebrafish retina anti-Rpe 65 in green and DAPI in blue, January 2023. URL https://zebrafishucl.org/antibodies-macdonald/rpe65.

85. John D. Fehilly, Tess McCann, Grace Ruddin, Joanna J. Kaylor, Hannah Grenville, Rebecca Ward, Kieran Wynne, Alicia Gómez Sánchez, Elin Strachan, Gabriel H. Travis, Ross F. Collery, and Breandán N. Kennedy. Germline disruption of retinal pigment epithelium-expressed zebrafish rlbp1b −/− results in selective dim light visual behavior deficits and provides a screening platform for evaluating the pathogenicity of human RLBP1 variants. The FASEB Journal, 39(12), June 2025. ISSN 1530-6860. doi: 10.1096/fj.202500600r.

86. Philip D. Kiser and Krzysztof Palczewski. Membrane-binding and enzymatic properties of RPE65. Progress in Retinal and Eye Research, 29(5):428–442, September 2010. ISSN 1350-9462. doi: 10.1016/j.preteyeres.2010.03.002.

87. Tünde Molnár, Oleg Yarishkin, Anthony Iuso, Peter Barabas, Bryan Jones, Robert E. Marc, Tam T. T. Phuong, and David Križaj. Store-operated calcium entry in müller glia is controlled by synergistic activation of TRPC and Orai channels. The Journal of Neuroscience, 36(11):3184–3198, March 2016. ISSN 1529-2401. doi: 10.1523/jneurosci.4069-15.2016.

88. H. Jingami, M. S. Brown, J. L. Goldstein, R. G. Anderson, and K. L. Luskey. Partial deletion of membrane-bound domain of 3-hydroxy-3-methylglutaryl coenzyme A reductase eliminates sterol-enhanced degradation and prevents formation of crystalloid endoplasmic reticulum. The Journal of cell biology, 104(6):1693–1704, June 1987. ISSN 1540-8140. doi: 10.1083/jcb.104.6.1693.

89. Nica Borgese, Maura Francolini, and Erik Snapp. Endoplasmic reticulum architecture: structures in flux. Current Opinion in Cell Biology, 18(4):358–364, August 2006. ISSN 0955-0674. doi: 10.1016/j.ceb.2006.06.008.

