## Supplementary figures and images for "A Müller glia-retinal pigment epithelium apical network surrounding cone outer segments in fish and mouse"

### FigS1

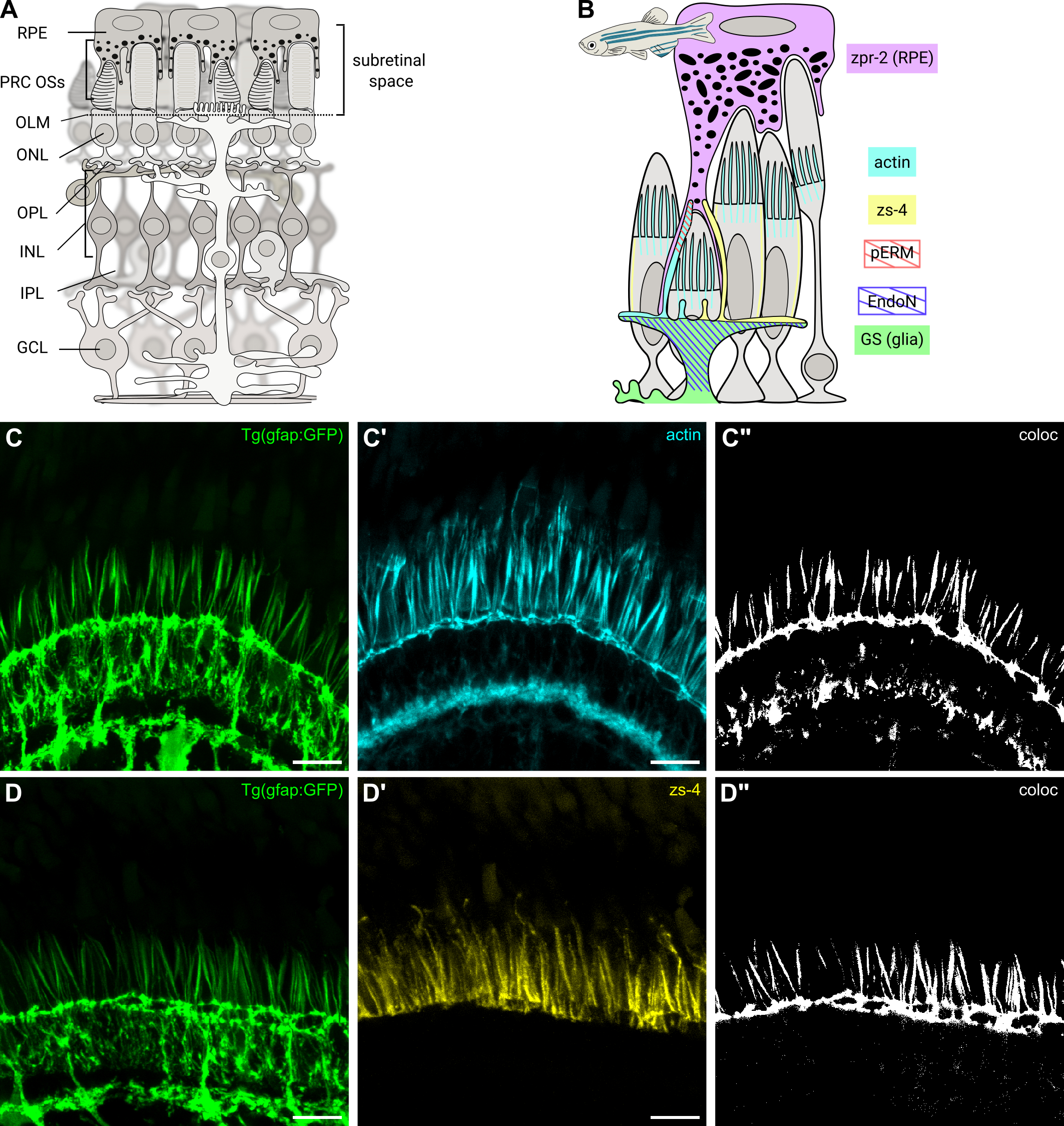

### FigS2

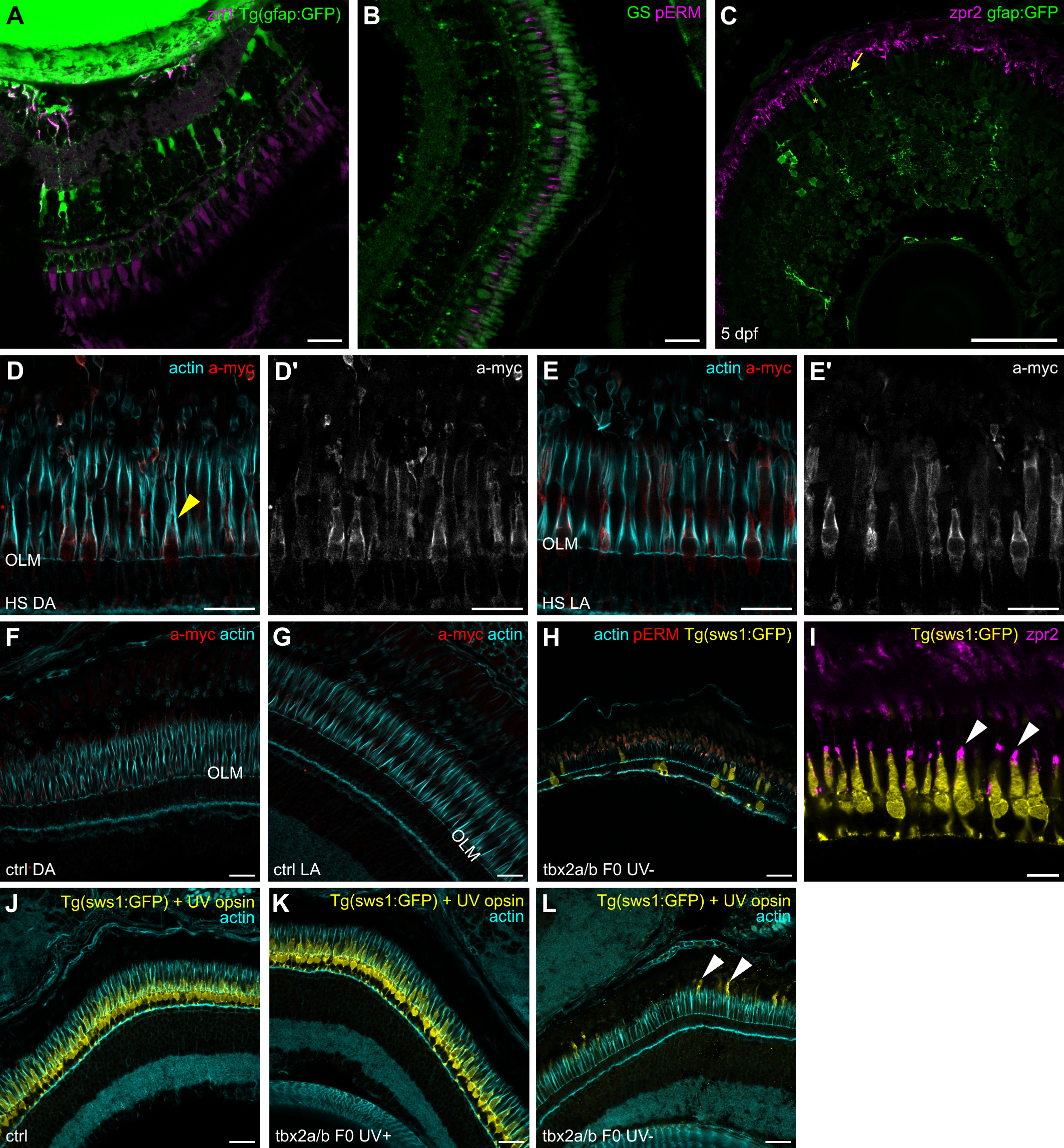

### FigS3

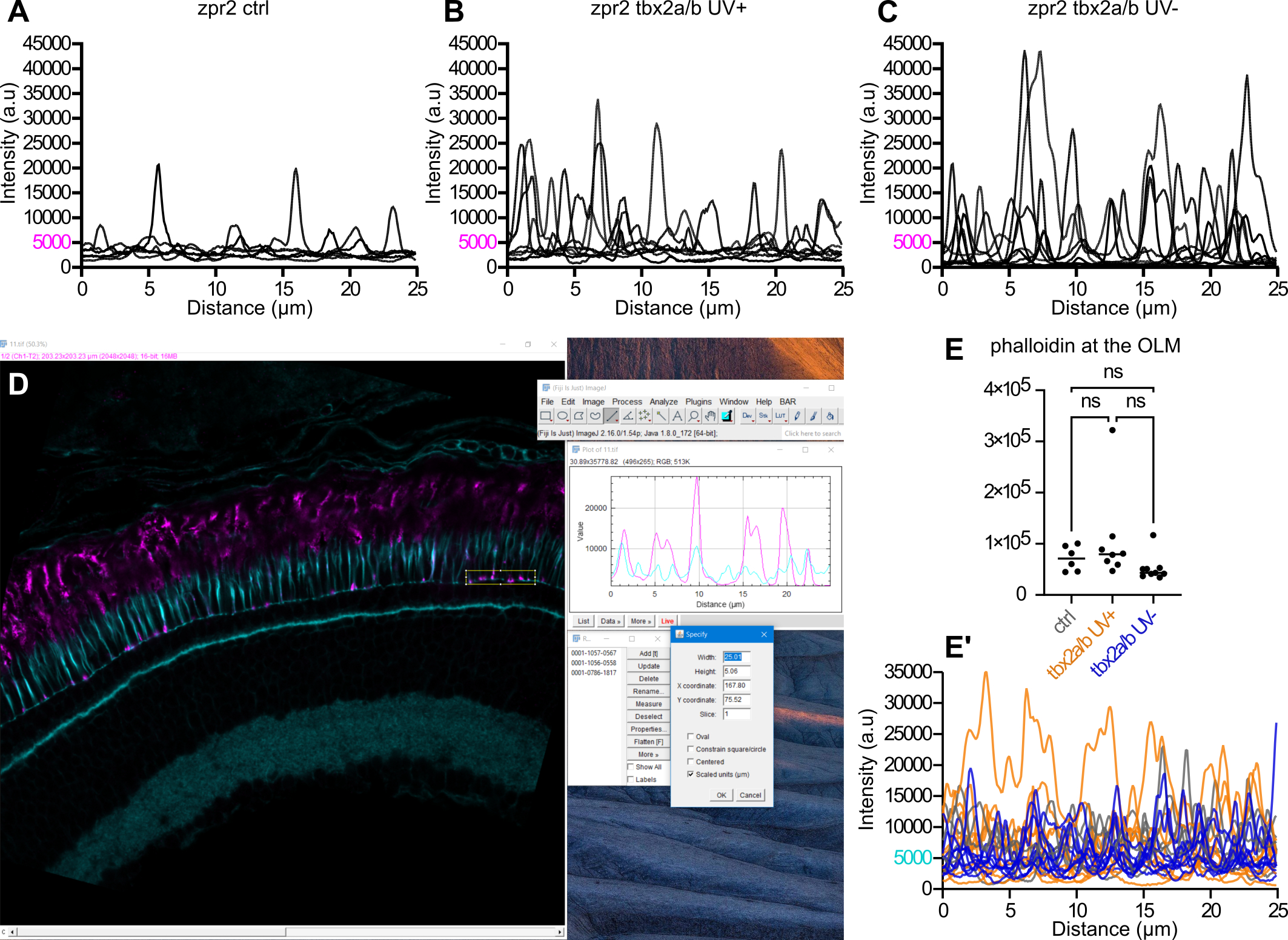

### FigS4

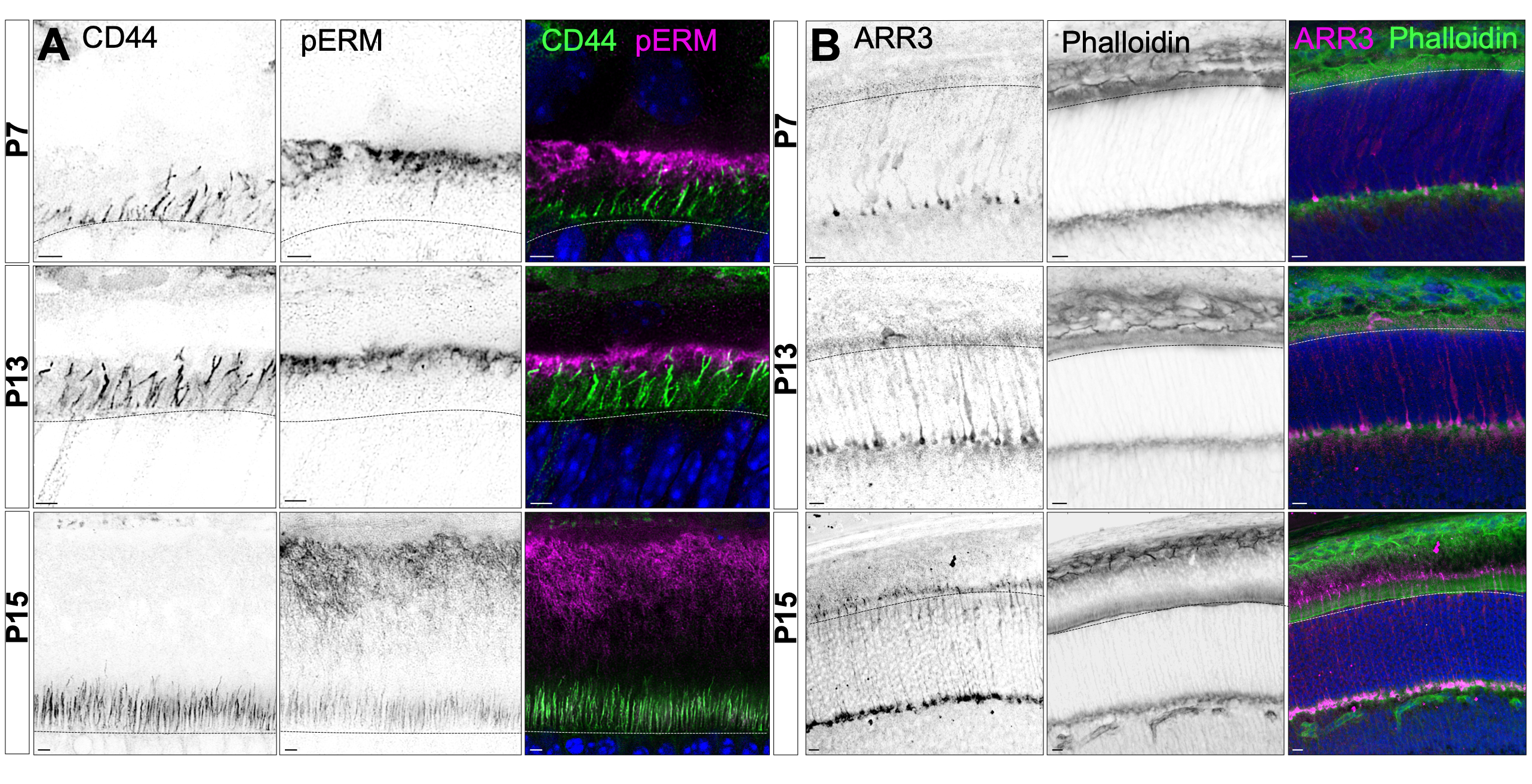

### FigS5

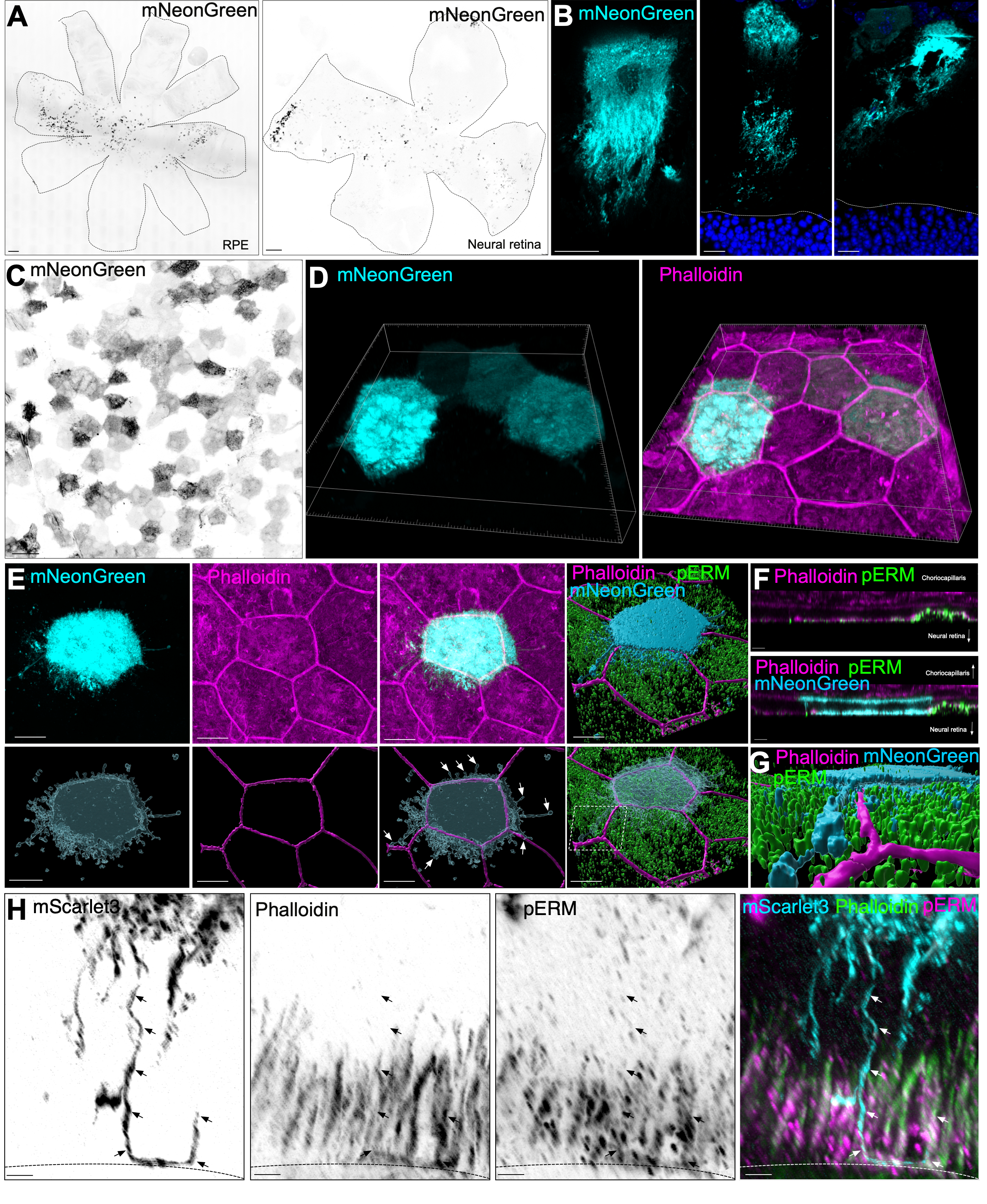

### FigS6

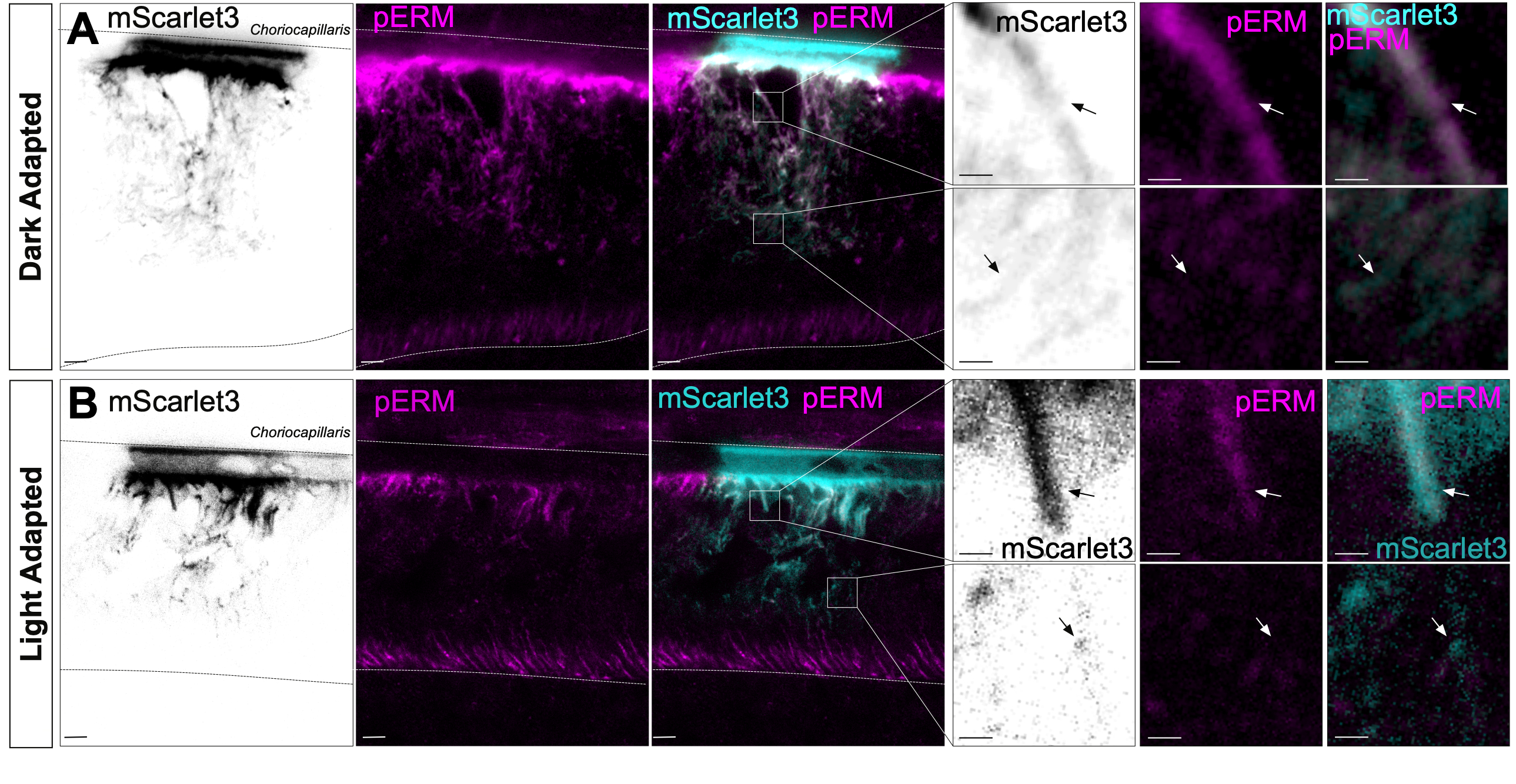

### FigS7

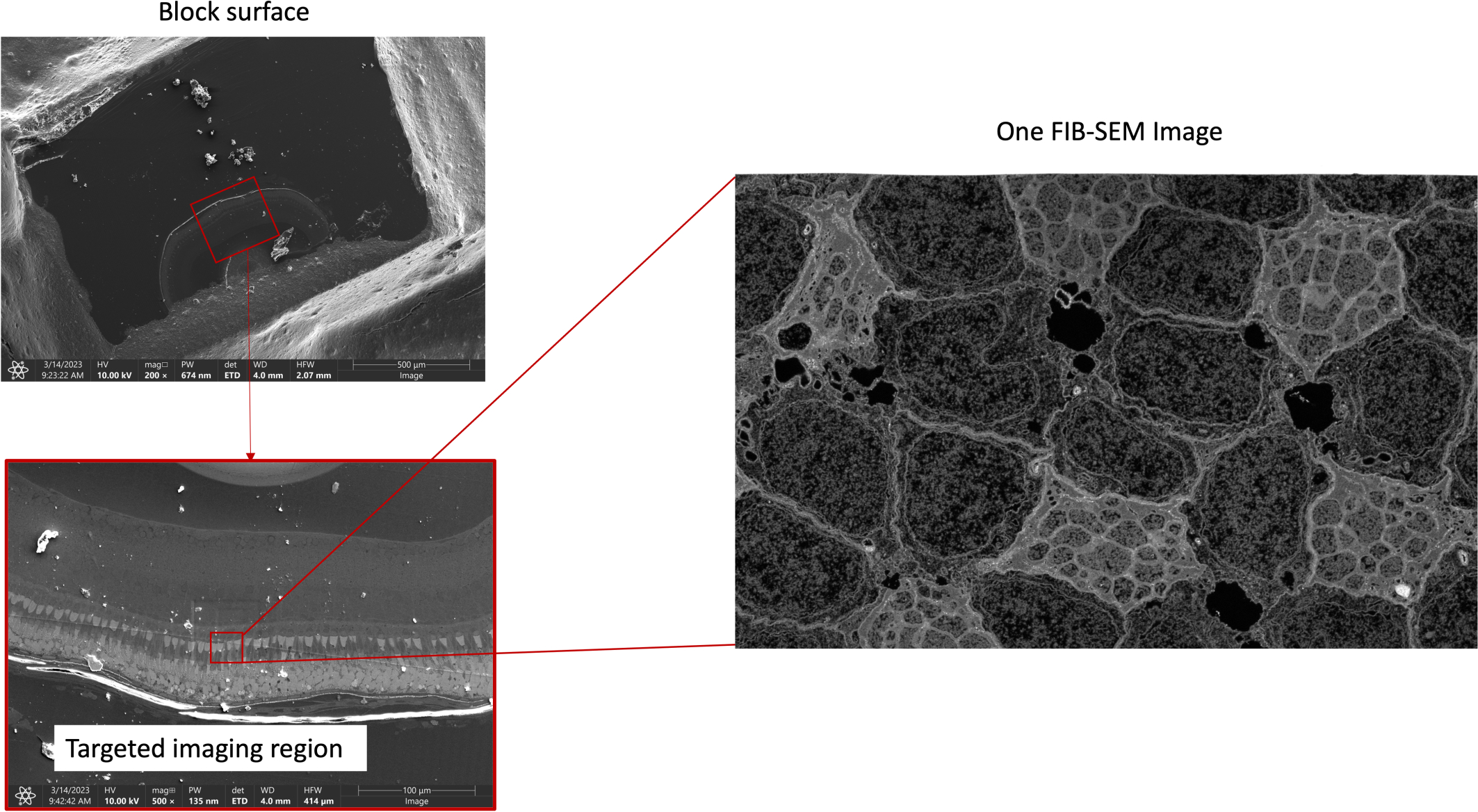

### FigS8

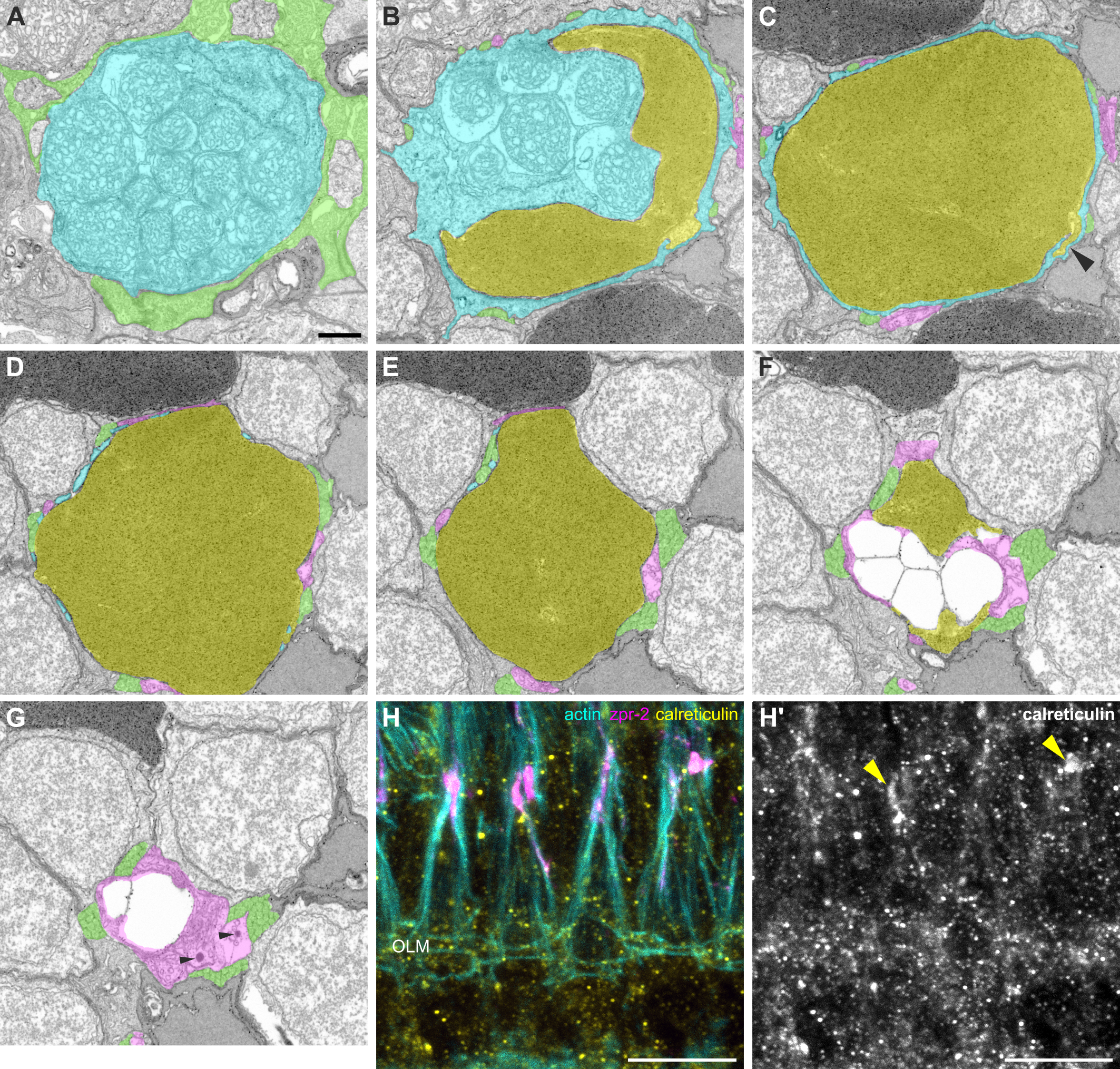

### FigS9

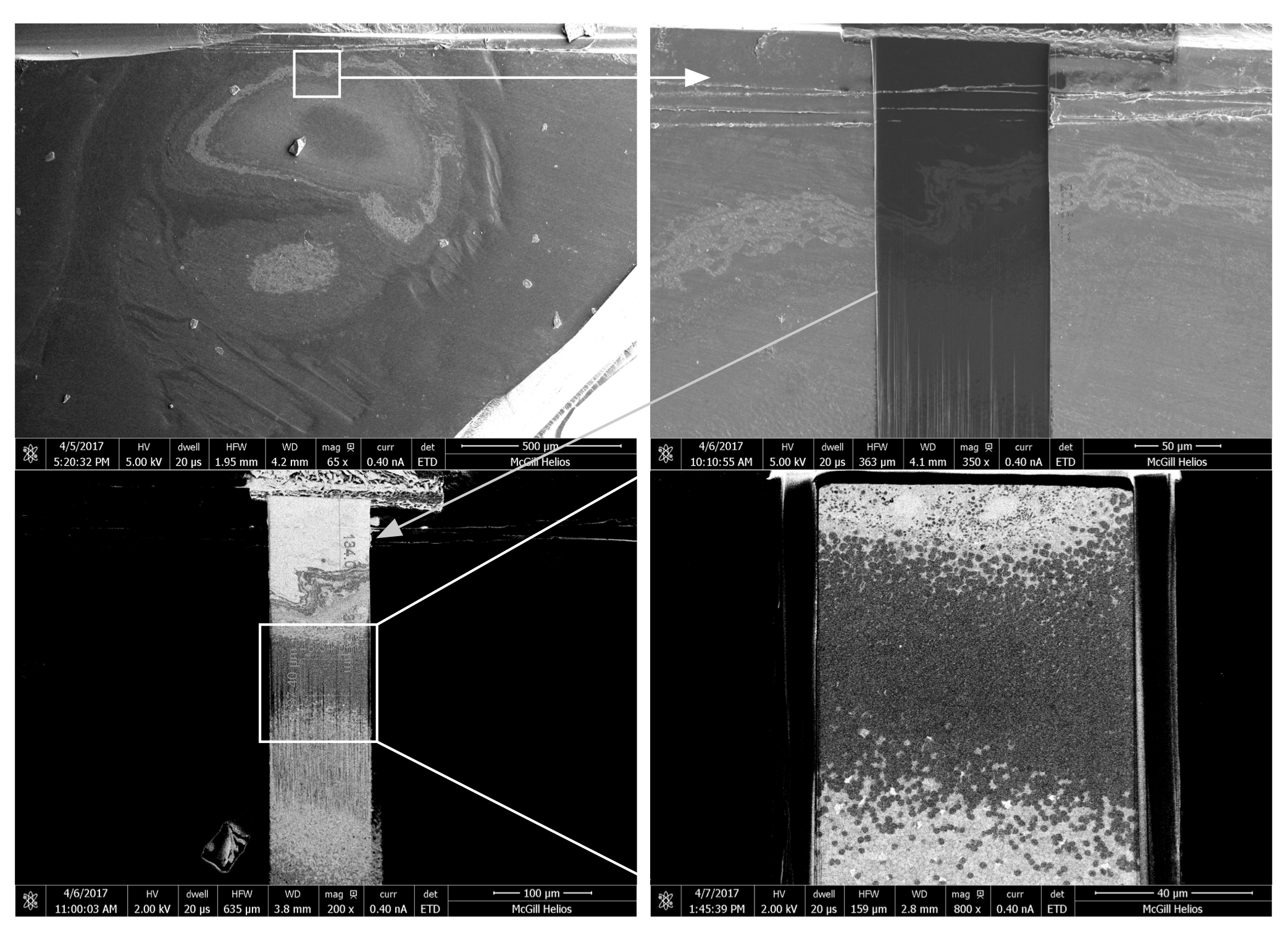

### FigS10

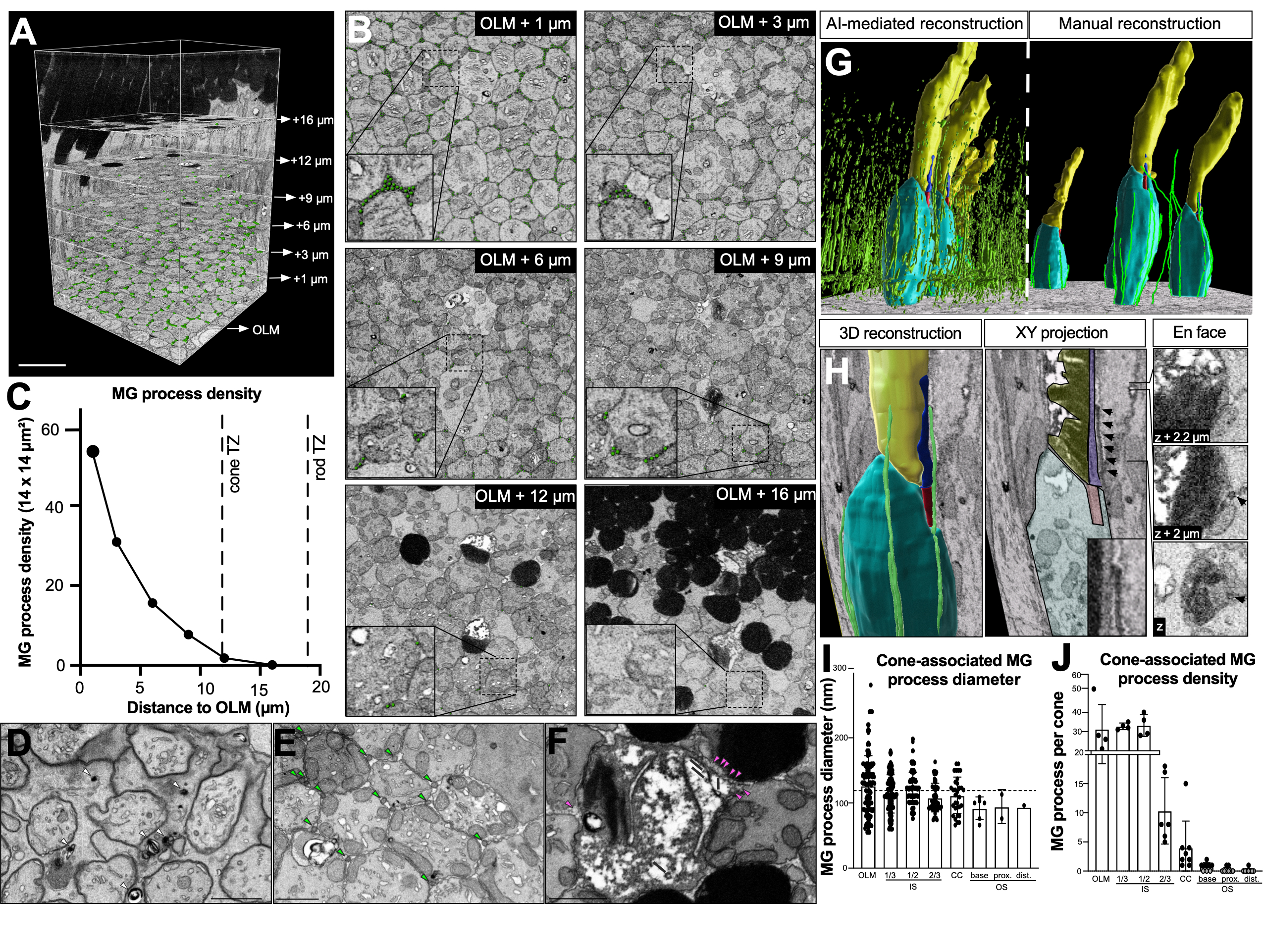
